# A 3D-printed multi-compartment organ-on-chip platform with a tubing-free pump models communication with the lymph node

**DOI:** 10.1101/2024.05.20.594865

**Authors:** S. R. Cook, A. G. Ball, A. Mohammad, R. R. Pompano

## Abstract

Multi-organ-on-chip systems (MOOCs) have the potential to mimic communication between organ systems and reveal mechanisms of health and disease. However, many existing MOOCs are challenging for non-experts to implement, due to complex tubing, electronics, or pump mechanisms. In addition, few MOOCs have incorporated immune organs such as the lymph node (LN), limiting their applicability to critical events such as vaccination. Here we developed a 3D-printed, user-friendly device and companion tubing-free impeller pump with the capacity to co-culture two or more tissue samples, including a LN, under a recirculating common media. Native tissue structure and immune function were incorporated by maintaining slices of murine LN tissue *ex vivo* in 3D-printed mesh supports for at least 24 hr. In a two-compartment model of a LN and an upstream injection site in mock tissue, vaccination of the multi-compartment chip was similar to *in vivo* vaccination in terms of locations of antigen accumulation and acute changes in activation markers and gene expression in the LN. We anticipate that in the future, this flexible platform will enable models of multi-organ immune responses throughout the body.

## INTRODUCTION

Molecular communication between organs is essential for life, both to maintain homeostasis and to respond rapidly to perturbation.^1^ A key example is vaccination, which requires proper drainage of signals from the site of injection to the draining lymph node (LN) to initiate early inflammatory activation and ultimately protective immunity. However, it is challenging to isolate the communication between specific organs using *in vivo* models because many organs contribute simultaneously through blood and lymphatic vasculature. Multi-organ-on-chip (MOOC) technology addresses this challenge by connecting compartmentalized models of select organs together under well-controlled conditions, often with circulating blood or lymph-like media.^2–14^ Given the critical role of vascular and interstitial flow rates *in vivo*,^2,15^ MOOCs must have precise and controllable flow speeds, including with thick 3D cultures. MOOCs with recirculating fluid flow provide biological feedback loops between organs, allowing accumulation of otherwise dilute secreted factors and minimal media consumption.^16^ However, despite their potential, MOOCs have not yet seen broad adoption by biomedical researchers.

For users focused on applications rather than microfluidics, the ease of use, compatibility with off-chip analysis, scalability for well powered hypothesis testing, and reproducible and affordable fabrication are all just as critical as biomimicry.^7,17^ These requirements are currently unmet for three reasons. First, most approaches for precisely controlled recirculating flow through MOOCs require extensive tubing and sophisticated electrical or pneumatic control systems, making them challenging for non-experts to implement.^18^ Closed-box commercial systems are beginning to be available and are designed to be easy to use, but at a high cost. Second, many OOCs house the cell or tissue culture permanently inside of a sealed device, such as on a membrane inside of a microchannel, making it difficult to add tissues on demand or to remove them for imaging, flow cytometry, or gene expression analysis during or after the experiment.^7^ Third, although many OOCs are hand-assembled in polydimethylsiloxane (PDMS), broad adoption for statistically powered biological experiments requires more reproducible fabrication methods, such as 3D printing, machining, or embossing.^19,20^ Thus, while there have been major advancements in MOOC technology, there is still a need for an easy-to-use multi-tissue chip that incorporates well-controlled fluid flow past biomimetic and experimentally accessible 3D cultures.

In addition to user-friendly technology, incorporation of organs of the immune system into MOOCs is an exciting frontier and is particularly dependent on inter-organ fluid flow. So far, MOOCs have primarily incorporated tissue-resident immune cells or recirculating white blood cells into existing models of lung, gut, brain, tumor, islets, etc.^3,21–24^ Incorporation of models of dedicated immune organs such as the lymph node (LN) is a major next step to model systemic immune responses such as vaccination, infection, and autoimmunity.^25,26^ LNs are small organs located along lymphatic vessels, where they filter flowing lymph fluid to detect and respond to pathogens (Fig. 1a).^27^ Vaccination takes advantage of this system by inducing the adaptive immune response to protect against infection, with initial responses occurring within hours due to fluidic transport from the injection site.^28^ Microscale and organoid models of the LN are still in early stages even in isolation,^25,29–35^ let alone in connection with other organs.^36–38^ The model with the longest history is *ex vivo* LN slice culture, which have been used to model the response to infection and vaccination in humans and animals for thirty years.^39–43^

**Figure 1.**
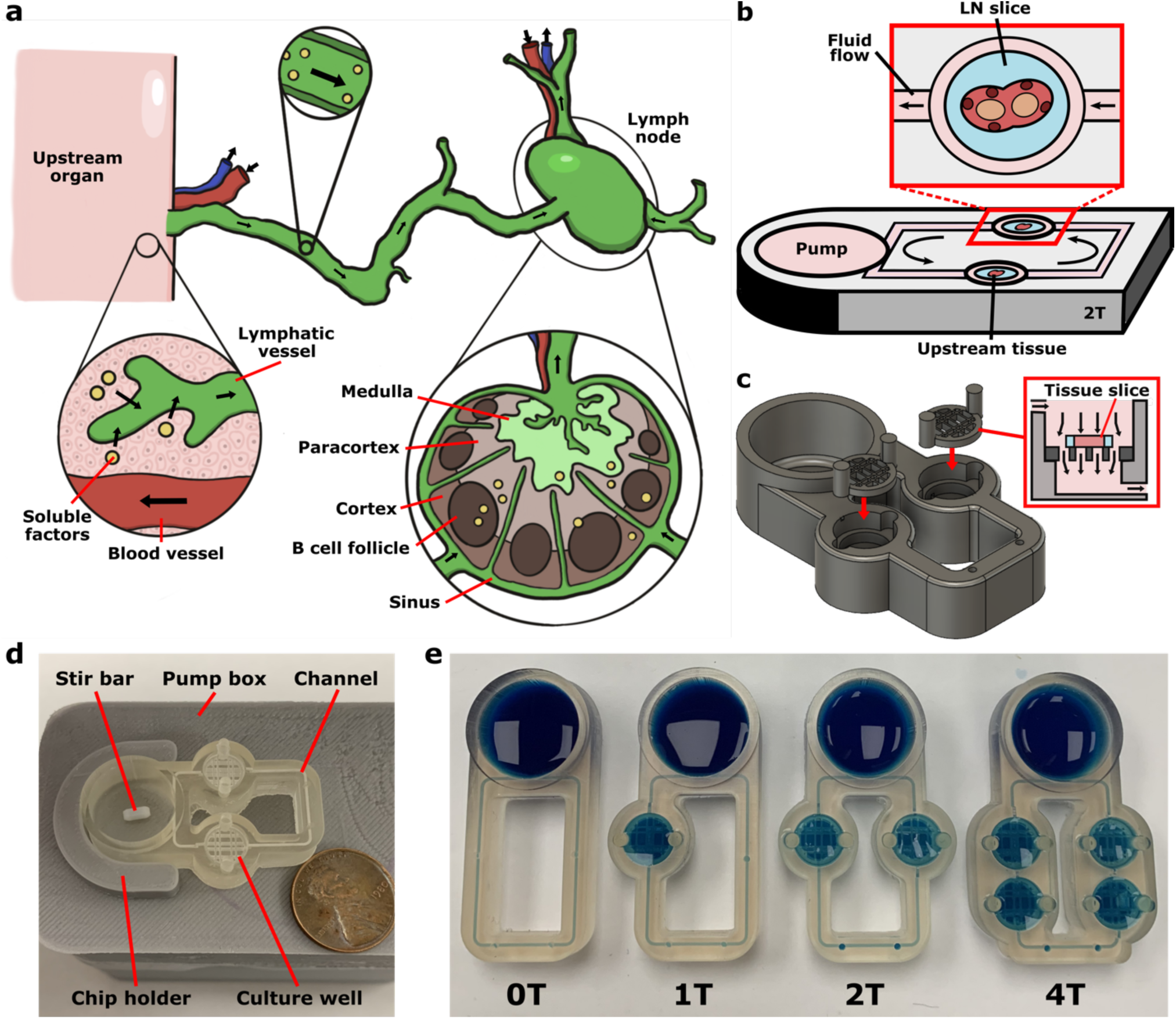
Modeling communication with the lymph node using a multi-compartment chip. (a) Illustration of communication via soluble factors (*yellow dots*) from an upstream organ to local lymph nodes via lymphatic vessels and interstitial fluid flow through each organ. (b) Schematic of the multi-compartment device, which consisted of a loop of channels containing wells for tissue slice culture connected to a pump well. (c) 3D rendering of the device showing the insertion of the removable mesh support into the open top of the culture well. The mesh support holds a tissue slice suspended within the well to enable flow perpendicular to the tissue. (d) Photo of a two-compartment device (ITX-PEGDA resin) mounted on the motor-based impeller pump external platform, with a US penny for scale. (e) Photo of four variations of the device (ITX-PEGDA resin) containing zero (0T), one (1T), two (2T), and four (4T) wells for tissue slice culture. Each chip was filled with blue food dye to visualize the channels.

Here, we developed a user-friendly, self-contained multi-compartment platform capable of multi-tissue co-culture under continuous recirculating fluid flow, and applied it to model the communication between the lymph node and upstream drainage sites. Inspired by prior work integrating pumps directly into a device to minimize tubing,^16,44–49^ and particularly by the simplicity of on-chip magnetic impeller pumps,^50–52^ we developed a 3D-printed multi-compartment device and companion tubing-free motor-based impeller pump. The system was designed to be customizable for the needs of the user, including to change the number of tissue compartments and volumes. As a proof-of-concept, we used this platform to develop a model of the acute response to vaccination in murine LN slices, and tested its fidelity against *in vivo* vaccination in terms of antigen drainage and processing, early markers of activation, and changes in gene expression.

## EXPERIMENTAL

### Device fabrication and assembly

The microfluidic devices and removable mesh supports were designed using Fusion 360. Designs for the devices and removable mesh insert were deposited in a public repository (see Data Availability below). Devices were printed in MiiCraft Clear resin (CADworks3D, Toronto, Canada) and in a custom PEGDA resin (ITX-PEGDA resin) formulated from a previously developed recipe.^53^ The PEGDA resin consisted of poly(ethylene glycol) diacrylate (PEGDA, 250 MW, Sigma Aldrich) as the monomer, phenylbis(2,4,6-trimethylbenzoyl)phosphine oxide (Irgacure 819, Sigma Aldrich) as the photoinitiator, and isopropylthioxanthone (ITX, Fisher Scientific, New Hampshire, USA) as the photoabsorber. Irgacure 819 and ITX were mixed with PEGDA (0.4% w/w) using a vortex mixer and dissolved for 30 min at 70°C.

Devices and mesh supports were printed using a CADWorks3D MiiCraft P110Y DLP printer (CADWorks3D, Toronto, Canada). All layers were printed at 50 µm at 100% power (5 mW/cm^2^), 1.25 s cure time, 4 s base cure time, 6 base layers, and 4 buffer layers. For post-processing, all printed parts were submerged in isopropyl alcohol (IPA) in a Form Wash (FormLabs, Massachusetts, USA) for 4 min, dried thoroughly with nitrogen, and placed in a Form Cure high-intensity UV light box (10 mW/cm^2^, FormLabs, Massachusetts, USA) for 1 min at room temperature. For device post-processing, channels were flushed with IPA using a wash bottle to clear channels of uncured resin before cleaning the chip in the Form Wash.

### Preparation of 3D-printed material for tissue culture

To improve biocompatibility of the 3D-printed material, the device and mesh supports were coated in Parylene-C as described previously.^54^ In brief, a film of ∼1 µm was achieved by adding 1.1 g of mixed isomers of Parylene-C (SCS, Inc. Indianapolis, IN, USA) to the Labcoater 2 parylene coater (SCS, Inc. Indianapolis, IN, USA) for gas-phase deposition onto the prints. Prior to culture of tissue in any 3D-printed device, the print and stir bar were sterilized by submerging in 70% ethanol for 5 min, followed by two 10 min rinses in 1xPBS (Lonza, Maryland, USA). Once rinsed, the materials were allowed to air dry for at least 30 min before use. The removable mesh supports were loaded into the devices as needed and filled with complete media to the specified volume per device. The chips were then loaded on to the external pump platforms, which were turned on and placed in the incubator for at least 30 min before use to reach 37 °C.

### Motor-based impeller pump assembly

The 3D-printed external housing, chip holder, and chip cover were designed using Fusion 360 and printed using 1.75 mm polylactic acid (PLA) filament (Flashforge, China) using a Monoprice Voxel 3D printer (Monoprice, California, USA). Designs for all of the pump-related files were deposited in a public repository (see Data Availability below). To assemble the motor-based impeller pump, a 6-12 V Mini DC motor (AUTOTOOLHOME) was inserted into the base of the printed housing. A custom magnet holder containing a small fan was 3D printed using the DLP printer described above in the ITX-PEGDA resin. The fan was included on the magnet mount to help push air down through the vents on the top of the motor and mitigate any heat buildup. Two 6-mm brushed nickel magnets (FINDMAG) with a strength of 0.008 T were glued using superglue into the magnet holder and mounted on the rotating pin of the DC motor.^51^ Each motor was connected to a mini digital DC voltmeter (2.5-30 V, MakerFocus, China) and a pulse-width modulation (PWM) low voltage DC potentiometer (ALDECO), both mounted to their respective holes within the housing base. Within each pump, three anodized aluminum heatsinks (1 g, Easycargo) were mounted along the sides to help distribute heat away from the DC motor. Once assembled, the housing top was initially glued together with hot glue, and the seam was sealed with an epoxy to generate a moisture-free environment within the box. The chip holder was glued to the top of the external housing centered over the DC motor. Each pump was connected to a 12 V DC female power connector (Chanzon), which was plugged into the 12 V AC DC power supply adapter wall plug (EWETON). A cord splitter was used to connect all 8 pumps to a single power supply. All wiring was connected using a tin-lead rosin-core solder wire (ICESPRING) and wrapped in heat shrink tubing (Eventronic, Germany).

A Teflon PTFE encapsulated magnetic stir bar, either 2 x 5 mm (VWR, Pennsylvania, USA) or 3 × 10 mm (Thomas Scientific, New Jersey, USA) were used as impellers. The 2 x 5 mm stir bar was used for all experiments unless noted otherwise. A digital laser photo tachometer (AGPtek, New York, USA) was used to measure the revolutions per minute (RPM) of the magnetic stir bar as it rotated. All RPMs reported were conducted for each individual pump for corresponding voltages, and are the average of three RPM measurements made at a consistent voltage. Stir bar stability and pump heat emission were measured as reported previously.^51^

### Characterization of experimental velocity within the device

The maximum velocity was measured experimentally as described previously.^51^ In brief, a drop of blue food coloring was pipetted into a port within the device and tracked using a Dino-Lite Edge 3.0 digital microscope (SunriseDino, California, USA). For each velocity measurement, we allowed for 1-2 min of equilibration after either changing the pump voltage or flushing the channels with a pipette. Images were collected over time with a timestamp to the millisecond decimal place, and the distance the dye moved over time was measured to determine fluid velocity.

### Animal model

All animal work was approved by the Institutional Animal Care and Use Committee at the University of Virginia under protocol #4042, and was conducted in compliance with guidelines from the University of Virginia Animal Care and Use Committee and the Office of Laboratory Animal Welfare at the National Institutes of Health (United States). Inguinal, axial, and brachial lymph nodes were harvested from female and male C57BL/6 mice (Jackson Laboratory, USA) under the age of 6 months following humane isoflurane anesthesia and cervical dislocation. There were no noticeable differences in viability (n = 2 M and 2 F), vaccine activation marker expression (n = 2 M and 1 F), or gene expression (n = 6 M and 6 F) between sexes, although these experiments were not powered to detect such differences. The lymph nodes were collected into “complete RPMI” media consisting of RPMI (Lonza, Maryland, USA) supplemented with 10% FBS (Corning, New York, USA), 1 x l-glutamine (Gibco Life Technologies, Maryland, USA), 50 U mL^−1^ Pen/Strep (Gibco Life Technologies, Maryland, USA), 50 µM beta-mercaptoethanol (Gibco Life Technologies, Maryland, USA), 1 mM sodium pyruvate (Hyclone, Utah, USA), 1x non-essential amino acids (Hyclone, Utah, USA), and 20 mM HEPES (VWR, Pennsylvania, USA).

### Preparation of lymph node slices

To generate lymph node slices, the inguinal, axial, and brachial lymph nodes were inserted into 6% w/v low melting point agarose (Lonza, Maryland, USA) in 1xPBS and punched into 5 mm blocks using a disposable biopsy punch (Royaltek).^43^ The 300 µm slices were generated using a Leica VT1000S vibratome (Illinois, USA) set to a speed of 90 (0.17 mm/s) and a frequency of 3 (30 Hz) while submerged in ice cold PBS. Slices were collected and placed in a 6-well plate containing ∼3 mL per well of complete media and placed in a sterile cell culture incubator (37 °C with 5% CO_2_) for 1 hr to rest prior to use.

### Measurement of viability of primary murine LN tissue

For resin cytotoxicity measurements, simple wells similar in size to the pump well were 3D printed and parylene coated as described above. The printed wells were inserted into a 12 well plate and filled with 1000 µL of fresh media, with empty well plate wells used as a plate control. The plate was equilibrated in the cell culture incubator at 37 °C for at least 30 min, after which LN slices from different nodes were randomly added to each well and cultured for 24 hrs.

For on-chip culture, the chips, mesh inserts, and stir bars were sterilized and dried as described above. Once dry, the mesh support(s) and stir bar were loaded into each device before filling with 1600 µL of fresh media. The channels were flushed through the ports using a pipette to ensure there were no bubbles hindering fluid flow. Chips were loaded onto the pump platforms and covered with a FDM 3D-printed cover, and the pumps were set to the required speed. The whole chip and pump assembly was equilibrated in the cell culture incubator for at least 30 min before tissue slices were added, after which slices from different nodes were randomly added to the culture wells containing mesh supports and cultured for 24 hrs. All cell and tissue viability experiments consisted of two identical experiments performed on different days pooled together to test reproducibility.

Following the culture period, LN slice viability was assessed using a CellTiter 96^®^ AQ_ueous_ One Solution Cell Proliferation Assay (MTS assay, Promega, Wisconsin, USA). Intact slices were added directly to 100 µL of media in a 96 well plate. A killed control was generated by adding 15 µL of 10X Lysis Buffer (CyQUANT LDH Cytotoxicity Assay Kit, Invitrogen, Massachusetts, USA) to the media for 30 min at 37°C. Next, 50 µL of fresh media was added to each well. Then, 30 µL of CellTiter One Solution Reagent was added to each well and incubated for 3 hrs at 37°C. At the end of the culture period, 100 µL of media was transferred to fresh wells on the same plate. Bubbles were removed by centrifugation for 5 min at 400 x *g*. Absorbance was measured at 490 nm using a CLARIOstar plate reader (BMG LabTech, Germany). The background absorbance from media-only controls were subtracted from the live and killed controls and samples.

### Soluble factor recirculation and capture on-chip

To generate mock tissue, biotinylated magnetic beads (0.5 µm beads, RayBiotech, Georgia, USA) were embedded in 3% w/v agarose (Lonza, Maryland, USA) in 1x PBS (Lonza, Maryland, USA), which was cast in a 3 mm punched hole within a 35 mm petri dish filled with solidified 6% w/v agarose. This was punched into 5 mm blocks with the bead-laden gel in the center using a disposable biopsy punch (Royaltek). Blocks were sliced to a thickness of 300 µm as described above using previously reported methods.^43^ Slices were collected and placed in a 6-well plate containing 1xPBS. Prior to protein insertion, the channels of the device (ITX-PEGDA resin) were blocked with BSA (bovine serum albumin) by filling the device 1% BSA (Fisher Scientific, New Hampshire, USA) in 1 x PBS (Lonza, Maryland, USA) and recirculating at 1250 RPM (1.7 V, 75 µm/s maximum channel velocity) for 1 hr. A biotin bead-loaded slice was added to a culture well within the device, and 5 µL of 200 µg/mL NeutrAvidin^TM^ Rhodamine Red^TM^-X (NRho, Fisher Scientific, New Hampshire, USA) was pipetted to the opposite culture well upstream of the slice. The biotin-bead loaded slices were removed from the device using the removable mesh insert and imaged at 0 hr (before NRho addition), 0.5 hr, 1 hr, 2 hr, 4.3 hr, and 24.6 hr using a Zeiss Axio Zoom macroscope (Carl Zeiss Microscopy, Germany). All images were analyzed in ImageJ, where the background-subtracted mean grey value of the entire tissue slice was calculated.^55^

### DQ-OVA capture in live LN slices on-chip

LNs were sliced as described above. To generate a killed control for on-chip and off-chip culture, during the 1 hr rest period after tissue slicing, a portion of the slices were transferred to a 24 well plate containing 2 mL of 35% ethanol for 30 min, then moved to a new well containing 1x PBS for a 30 min rinse. Live and killed slices were cultured both on-chip and off-chip in the presence of DQ-ovalbumin (DQ-OVA, Invitrogen, Massachusetts, UVA). Following the 1 hr rest/kill periods, the slices were added to either a well plate or the downstream tissue culture well of a 2T chip, both filled with 1800 µL of complete media. For the on-chip conditions, 10 µL of 500 µg/mL DQ-OVA was added to the upstream culture well. Similarly, 10 µL of 500 µg/mL DQ-OVA was added directly to each off-chip culture well of the well plate. The slices were imaged at 0 hr (before DQ-OVA addition), 2 hr, 4 hr, and 24 hr on the Zeiss Axio Zoom macroscope. The off-chip slices were imaged within the well plate, while the on-chip slices were removed on the mesh supports and placed on a sterile petri dish for imaging before re-insertion. The slices were analyzed using ImageJ, where the background-subtracted mean grey value of the entire tissue slice was calculated. All images were leveled the same unless stated otherwise. The data presented consisted of two identical experiments performed on different days to demonstrate reproducibility.

### Comparative vaccination on-chip, *in vivo*, and well plate

For *in vivo* vaccinations, male and female C57Bl/6 mice were vaccinated subcutaneously with four 50 µL injections per animal at the shoulders and hips. The vaccine consisted of 100 μg/mL R848 (InvivoGen) and 500 μg/mL rhodamine-labeled ovalbumin (Rho-OVA) in sterile 1x PBS, or PBS as a vehicle control. The skin-draining lymph nodes (axillary, brachial, and inguinal) were harvested either 24 hrs later for the imaging experiment or 6 hrs later for the RNA sequencing experiment.

For on-chip and well plate (wells) conditions, “mock skin” blocks were generated by punching 5 mm cylinders out of 6% agarose gel with a biopsy punch. The cylinders were placed on parafilm in a petri dish and excess liquid was gently removed. The vaccine solution, either 0.2 µg/mL R848 and 2 µg/mL Rho-OVA or PBS, was pipetted directly on top of each gel block and allowed to passively diffuse into the gel for 30 min. In parallel, LN slices were collected from naive animals and sliced as described above. For the on-chip condition, the LN slice was added to the filled 2T device in the downstream well, and the mock skin was loaded on to a removable mesh support and loaded on-chip in the upstream well. For the well plate condition, the LN slice was added to a media-filled well in a 12-well plate, followed by the mock skin next to it. After 6 hours, the LN slices were collected for both the imaging experiment and the RNA sequencing experiment.

### Immunostaining and confocal fluorescence microscopy for vaccination experiment

Upon collection, all slices were immunostained as described previously.^56^ Briefly, slices were blocked with anti-mouse CD16/32 for 30 min in a cell culture incubator. An antibody cocktail containing: BV421 CD86, AF488 CD69, AF647 CD40, and Starbright Violet 670 CD19, was added for 1 hr (Table S1). Prior to imaging, the slices were washed for 30 min in 1x PBS. Confocal microscopy was performed on a Nikon A1Rsi confocal upright microscope using 405, 487, 561 and 638 nm lasers paired with 450/50, 525/50, 525/50 and 685/70 nm PMTs on a GaAsP detector, respectively. Starbright Violet 670 was excited off the 405 nm laser and detected with the 685/70 PMT. Images were collected with a 40×/0.45NA Plan Apo NIR WD objective.

Images were analyzed using Image J (version: vt1. 53t). Three regions of interest (ROIs), the entire slice, CD19+, and CD19-region, were defined by thresholding the outline of the tissue or the CD19 signal in the Cy5 channel. In each ROI, the mean grey value (MGV) of CD86, CD69 and CD40 were quantified. Each image was corrected for spillover from other channels by subtracting average MGV from three fluorescent minus one (FMO) images. The reported results are pooled from three identical experiments performed on different days.

### RNA isolation and sequencing

RNA was isolated from intact LN tissue and slices using the RNeasy Plus Mini (Qiagen, Germany). Samples were kept on ice throughout isolation process and surfaces were cleaned using RNase AWAY (Molecular BioProducts, California, USA). For both R848 + Rho-OVA and PBS *in vivo* conditions, each sample consisted of 2 intact LNs from a single mouse. For both R848 + Rho-OVA and PBS well plate and on-chip conditions, each sample consisted of 3 pooled LN slices from a single mouse. A sample key can be found in Table S2.

Intact LNs or LN slices removed from agarose were placed in 350 µL of RLT buffer with 1% 14.3 M ß-mercaptoethanol (Sigma-Aldrich, Missouri, USA) and homogenized using a Model 150 VT Ultrasonic Homogenizer (BioLogics, Inc, North Carolina, USA) at 30% power for 7 pulses over 2 min, then vortexed at maximum speed for 1 min. The samples were centrifuged for 3 min at maximum speed. The supernatant was transferred to a gDNA Eliminator spin column in a 2 mL collection tube and centrifuged for 30 s at 8000 x g to remove DNA from the sample. The column was discarded, and 350 µL of 70% ethanol was added to the flow-through and mixed. The 700 µL of sample was added to a RNeasy spin column placed in a 2 mL collection tube and centrifuged for 15 s at 8000 x g and the flow-through was discarded. Next, 700 µL of RW1 buffer was added to the spin column and centrifuged for 15 s at 8000 x g and the flow-through was discarded. 500 µL of RPE buffer was added to the spin column and centrifuged for 15 s at 8000 x g and the flow-through was discarded. Another 500 µL of RPE buffer was added to the spin column and centrifuged for 2 min at 8000 x g. The spin column was placed in a new 2 mL collection tube and centrifuged for 1 min at full speed to eliminate residual buffer. To elute the RNA, the spin column was placed in a new 1.5 mL collection tube and 50 µL of RNase-free water was added directly to the spin column membrane. The spin column and 1.5 mL tube were centrifuged for 1 min at 8000 x g to elute the RNA.

Following RNA isolation, the sample purity and concentration were measured using a ND-1000 NanoDrop spectrophotometer (Thermo Fisher Scientific, Massachusetts, USA). All samples per condition from a single experiment were pooled to improve RNA yield, where two samples from different mice were pooled for every experiment. After pooling, six samples remained (*in vivo* R848 + Rho-OVA, *in vivo* PBS, well plate R848 + Rho-OVA, well plate PBS, on-chip R848 + Rho-OVA, and on-chip PBS), with each sample containing RNA from multiple experimental and biological replicates (Table S5). The experiment was run in triplicate and all samples were stored at −80 C until sequenced. The RNA samples were submitted to NovoGene Co (California, USA) for sequencing. Following quality control, a non-directional low-input eukaryotic mRNA library was prepared. Bulk RNA sequencing was performed using a NovaSeq PE150 with 6 G of raw data per sample.

### Bioinformatics analysis

The RNA-seq data analysis was done by the University of Virginia Bioinformatics Core (RRID: SCR_012718). On average, 22 million (rages from 21 to 27 million) paired end reads (150 bases long read) were received for each of the replicates sufficient for gene level quantitation. The read quality was assessed using the fastqc program (http://www.bioinformatics.babraham.ac.uk/projects/fastqc/), and raw data quality report was generated using the MultiQC tool.^57^ Adaptor contamination observed in few samples were removed using “cutadapt” program.^58^ The “splice aware” aligner ‘STAR’ aligner was used for mapping the reads.^59^ Prior to mapping, a mouse reference index was constructed based on the GRCm38 mouse genome reference (Mus_musculus.GRCm38.dna.primary_assembly.fa & Mus_musculus.GRCm38.91.chr.gtf), and the sjdbOverhang parameter was set to 149 to match the read length of the samples. Subsequently, read mapping and quantification were conducted; more than 95% of the reads mapped to mouse genome and transcriptome.

Gene-based read counts were derived from the aligned reads, and subsequently, a count matrix was generated, serving as the input file for the analysis of differential gene expression. The DESeq2 package was used to conduct the differential gene expression analysis.^60^ Low expressed genes (genes expressed only in a few replicates and with low counts) was excluded from the analysis before identifying differentially expressed genes. Data normalization, dispersion estimates, and model fitting (negative binomial) were carried out with the DESeq function.

The log-transformed, normalized gene expression of 500 most variable genes was used to perform an unsupervised principal component analysis. The differentially expressed genes were ranked based on the log 2-fold change and FDR corrected p-values. Principal component analysis, z score heatmap, and gene set enrichment analysis (GSEA) for Hallmark pathways were carried out in R with the following three comparisons: 1) *in vivo* R848 + Rho-OVA / *in vivo* PBS, 2) on-chip R848 + Rho-OVA / on-chip PBS, and 3) wells R848 + Rho-OVA / wells PBS. Pathway analysis was performed using fgsea package in Bioconductor R (https://bioconductor.org/packages/release/bioc/html/fgsea.html). The reference database for mouse pathway enrichment analysis comprised Hallmark gene sets from the msigdb.^61–63^

## RESULTS AND DISCUSSION

### Customizable multi-compartment 3D-printed platform

The major design goals when designing the multi-compartment chip were 1) fast and reproducible fabrication; 2) expandable design to accommodate the co-culture of two or more tissues; 3) easy tissue insertion and removal for timecourse imaging without tissue damage; 4) biocompatibility with tissue slice culture; and 5) recirculating fluid flow on-chip. Inspired by the principles of our previous PDMS prototype,^37^ we developed a monolithic 3D-printed device that consisted of a loop of channels that connected varying numbers of tissue culture wells. The culture wells were in line with a pump well^51^ for recirculation of media and secreted molecular cues (Fig. 1b-d). The use of resin 3D printing provided a semi-transparent, easily customizable device. This fabrication method enabled complex 3D architectures such as sloped channels and mesh supports for slice culture, which would be challenging to produce using traditional soft lithography fabrication.^51,64^ We designed a series of monolithic devices with zero to four culture wells (0T – 4T) to illustrate the flexibility of the platform while retaining its simplicity (Fig, 1e), each printing in <1 hr. Users may select the device that is best suited for their specific application, based on the intended number of culture chambers. The multi-compartment chips had 500 µm channels for reproducible fabrication on the printer used; in the future, channels could be made smaller by using different liquid resin formulations and higher resolution printers.^53,65–68^

In principle, this system is compatible with 2D, 3D, or explant cultures inserted into the culture wells. Here, we incorporated live LN tissue slices to maintain the spatiotemporal organization of this organ and make it accessible for imaging and stimulation.^37,43,69^ Tissue slices are often cultured using perfusion to increase nutrient and gas exchange, and slices have a long history of incorporation into microscale perfusion devices.^37,38,69–78^ Here, to allow the fragile slices to be easily added on demand and removed repeatedly for timecourse imaging, slices were placed on a removable mesh support (Fig. 1c, Fig. S1), which could be quickly removed from the device using standard tweezers. Unlike commercially available transwell inserts with tall plastic walls, the 3D printed removable mesh was of a depth to hold the tissue between the channel inlet at the top of the chamber and outlet at the bottom of the chamber, thus providing transverse flow to carry signals downstream. We incorporated gaps in the mesh support around the edge of the tissue slice to limit the resistance through the fluidic loop.^37^

### Compact, tubing-free impeller pump and motor-based pump platform

Most standard pumps and pneumatic pressure controllers are bulky or incompatible with cell culture incubators, and thus require long tubing that is prone to bubbles, contamination, or disconnection. To avoid these issues, we developed a companion impeller pump with the following major design criteria: 1) no tubing, few wires, and simple controls for ease of use; 2) tunable recirculating fluid flow at biomimetic rates; 3) minimal media volume to reduce dilution; 4) low cost; 5) small footprint; and 6) low heat output. To achieve the first two criteria, the tubing-free impeller pump was driven by magnets rotated by a small DC motor inside a custom electronic control box totaling to ∼$35 (Fig. 2a,b). The pumps fit eight per row (Fig. 2c) for a theoretical total of 48 pumps per incubator. The spinning magnets drove the rotation of a magnetic impeller on-chip to generate recirculating fluid flow in the connecting channel loop (Fig. 2d, Movie S1). For simplicity, here we used commercially available stir bars of different sizes as impellers; in past work, we have also used 3D-printed cross-shaped impellers^51^ and vaned impellers successfully (unpublished results). With the motor and stir bar, the fluid volume in the chip was reduced by > 2-fold and the footprint of the pump by 2-fold compared to an earlier prototype.^51^ We confirmed that the stir bar revolutions per minute (RPM) increased linearly with voltage (Fig. S2a) and was stable for 90 hrs (Fig. S2b).^51^ In a 10-day test of heat output by 8 pumps, temperatures in the incubator remained within the acceptable range (+/− 1 deg) (Fig. S2c).^79^

**Figure 2.**
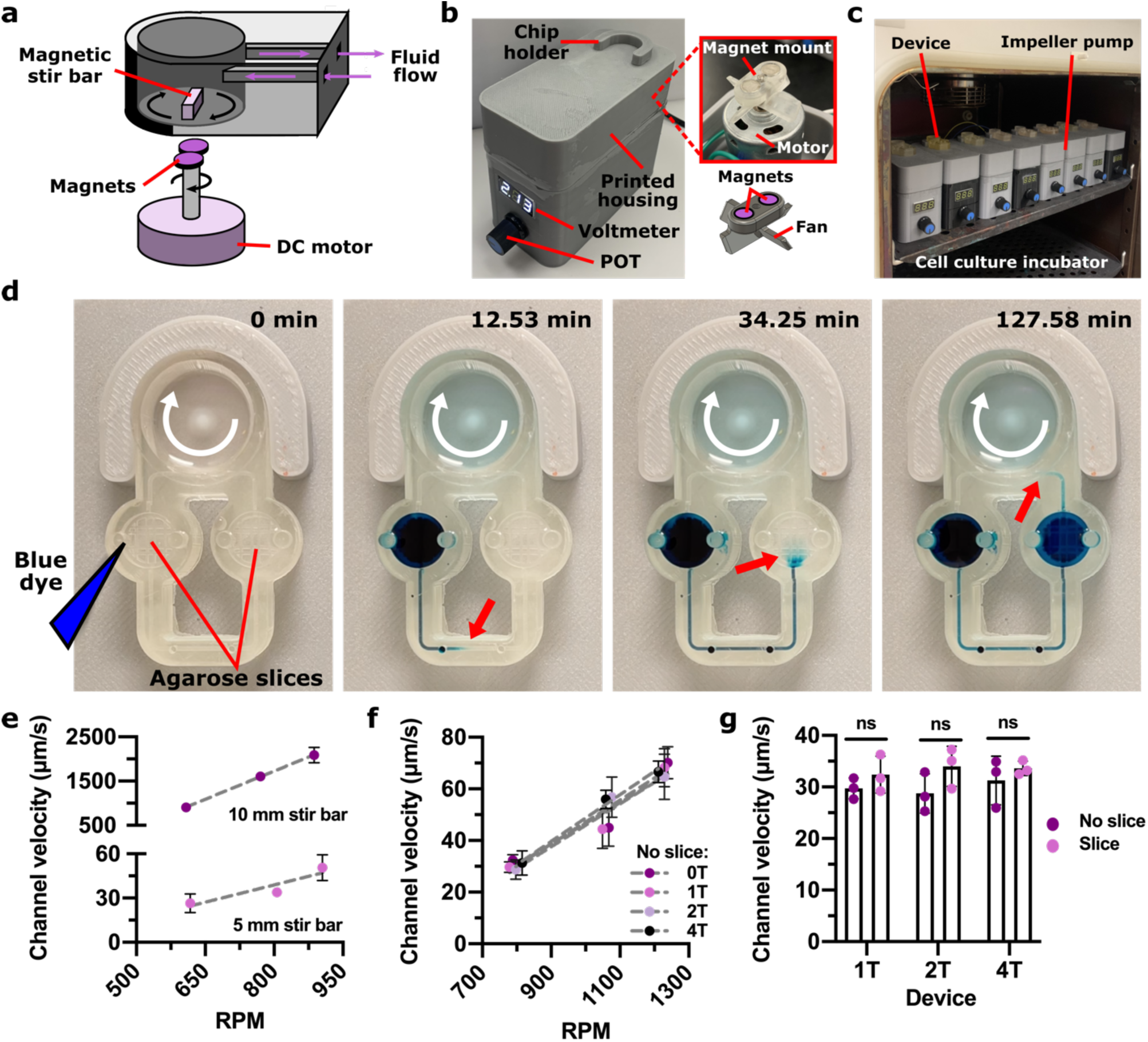
Motor-based impeller pump for fluid flow recirculation and control on-chip. (a) Schematic of the approach for impeller rotation. (b) Photos of the pump platform showing the outside of the pump box and (*right*) the interior of the pump. (c) Photo of eight impeller pumps on a shelf in a standard cell culture incubator, each holding a single multi-compartment chip. (d) Time-lapse images of recirculating fluid flow on a 2T device (Clear resin) with an agarose slice in each well (5 mm stir bar, 1000 RPM). Blue dye was inserted in the upstream culture well, and over time, moved through the channel to the downstream culture well and then the pump well (dye front marked with red arrow). The stir bar rotated clockwise within the pump well (white arrow). (e-f) Experimentally measured maximum velocity within the channel in (e) a 0T device (no culture wells) using a 10 mm stir bar and a 5 mm stir bar, and (f) in varied device designs (0T – 4T) with a 5 mm stir bar, all with no slice added. Dots and error bars represent mean and standard deviation; some error bars too small to see. (g) Experimentally measured maximum velocity in the channel at a low RPM (1.2 V, 780 RPM; 5 mm stir bar), with and without the addition of an agarose slice in each device. Each dot represents one velocity measurement. Results were compared using an unpaired t test (n = 3). ns indicates p > 0.1.

### On-chip fluid recirculation at controllable speeds

We made use of this pump to recirculate media and secreted factors through the chip for communication between culture compartments. As future applications may also include modeling of transport through vasculature, we tested how well the chip and pump achieved a range of physiologically relevant flow regimes, ranging from µm/s to mm/s. Whereas we previously showed that channel velocity was controlled by the physical geometry of the chip (e.g. channel width and length and pump well geometry), here we quantified the velocity through a fixed chip geometry as a function of user-controllable parameters.^51^ Doubling the length of the stir bar from 5 to 10 mm increased the channel velocity from tens of µm/s to mm/s, respectively, thus providing access to different flow regimes (Fig. 2e). Varying the volume of fluid added to the device also tuned the channel velocity, with reduced speeds as the meniscus rose away from the entry point of the channel (Fig. S3a,b). On the other hand, channel velocities were comparable between each device variation (0T, 1T, 2T, and 4T) (Fig. 2f) and with and without agarose slices added (Fig. 2g). Thus, at least with the current mesh support, the culture wells and mock tissue slices did not significantly increase the resistance of the microfluidic loop, providing robust flow control through the channels at wide ranges of fluid flow regimes.

### Tissue permeability and channel velocity control interstitial fluid speeds and soluble factor delivery in 3D computational model

Next, we determined the predicted range and spatial distribution of interstitial flow rates and mass transport through the tissue. Interstitial flow rates *in vivo* are widely variable, with typical estimates ranging from 0.01 – 10 µm/s.^80–82^ In a pressure-driven system, the hydraulic permeability of the tissue has a major impact on expected interstitial flow rate.^30,81^ To capture these effects, we developed a 3D finite element model of the tissue in the culture well using COMSOL Multiphysics (Fig. 3a,b, see ESI Methods). The simulated tissue and agarose were placed atop an impermeable mesh support and modeled as porous matrices of defined permeability and porosity. The permeability of the lymph node will significantly impact the interstitial fluid flow and molecular transport in the tissue, but this feature is largely unmeasured. Here, we tested the range of available predictions for LN permeability from 10^−10^ to 10^−12^ m^2^.^30,83^

**Figure 3.**
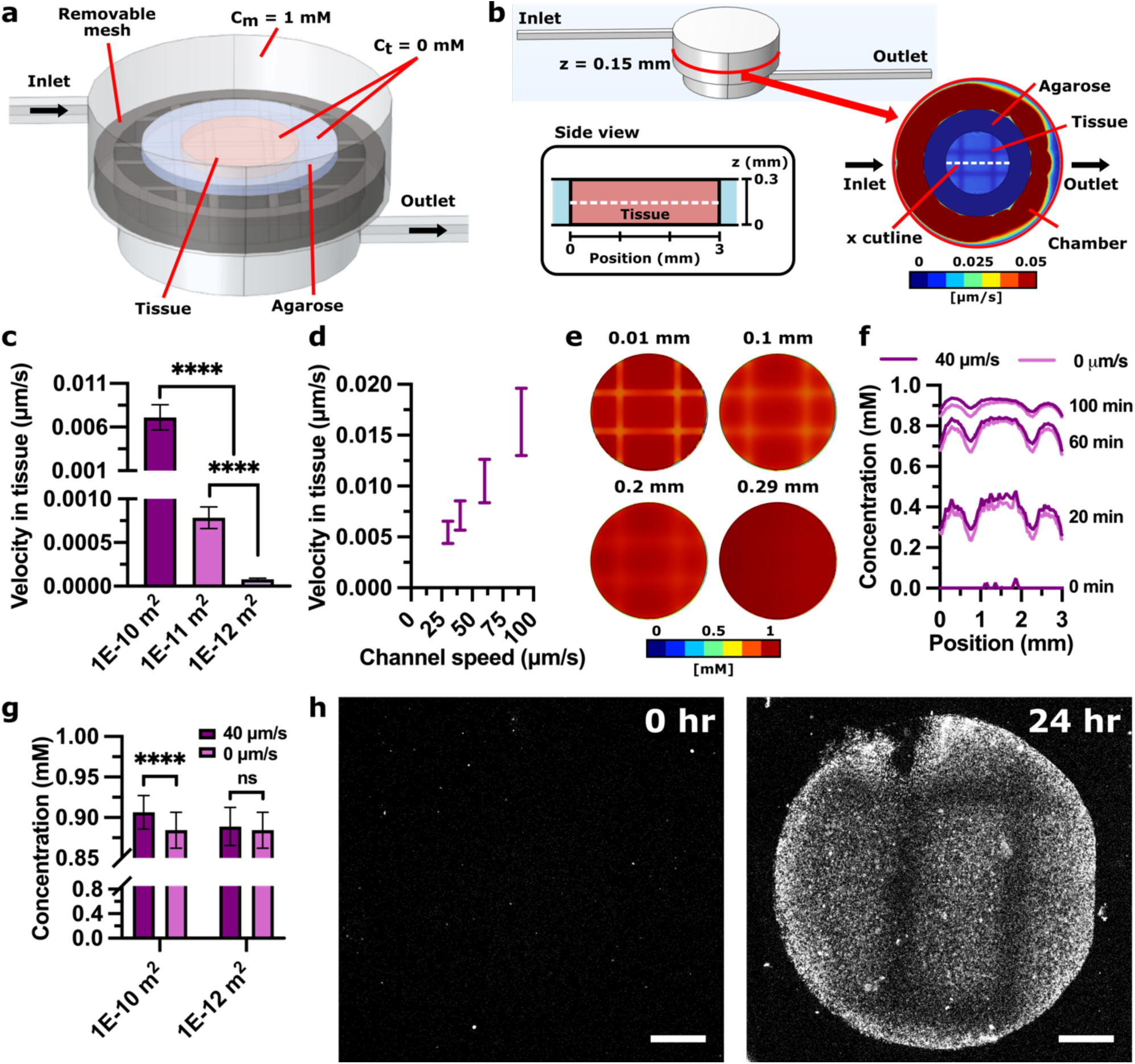
Simulated fluid velocity and tracer concentration through a tissue slice. (a) Geometry of the 3D COMSOL simulation, colored to show the tissue (*pink*) embedded in agarose (*blue*) resting on the removable mesh support (*dark grey*). The inlet channel intersects the top of the culture well, while the outlet channel intersects the bottom on the opposite side. (b) The velocity and tracer concentration were measured along cutlines along the x axis in line with the inlet and outlet channels, at different z planes. (c-d) Predicted velocity in tissue for (c) a range of tissue permeabilities at an inlet speed of 30 µm/s and (d) a range of inlet speeds at a permeability of 1×10^−10^ m^2^. One-way ANOVA with Tukey post-hoc tests. **** indicates p < 0.0001. In d, the error bars represent the range of velocity along the cutline. (e) Color plots of protein concentration in the tissue domain at 100 min at a permeability of 1×10^−10^ m^2^, showing different z planes (mm). (f) Predicted protein concentration in tissue over time with flow (40 µm/s) and without flow (0 µm/s) at a tissue permeability of 1×10^−10^ m^2^ on a central cutline (z = 0.15 mm) (g) Predicted protein concentration in tissue as a function of tissue permeability and flow on a central cutline (z = 0.15 mm) at 100 min. Bars represent standard deviation. One-way ANOVA with Tukey post-hoc tests. **** indicates p < 0.0001, ns indicates p > 0.2. (h) Representative images of the effect of the mesh support, with NRho (white) captured within the biotin region of the agarose slices at 0 hrs and 24 hrs at a channel speed of 30 µm/s. Scale bar shows 500 µm.

With the current mesh geometry, we anticipated that most of the fluid would pass through the gaps in the mesh support rather than through the tissue (illustrated in Fig. 1c). Indeed, only when the simulated tissue was more permeable (1×10^−10^ m^2^) did the mean interstitial flow velocities overlap with the lower bound of physiological interstitial fluid flow, ranging from 0.006 – 0.02 µm/s as a linear function of channel speed (Fig. 3c,d; Fig. S4a). Tissues with lower permeabilities had negligible interstitial fluid flow. The predicted shear stress in the tissue and the channels was lower than physiological shear stress in all cases (Fig. S5a,b), and thus was not expected to impact tissue viability significantly. In the future, it may be possible to increase the velocity and shear stress within the tissue to within physiologically relevant ranges by reducing the open space around the mesh, increasing the channel speed, or increasing the viscosity of the cell culture medium.

As one of the key functions of a multi-compartment chip is to transport molecular cues from the media into each tissue, we assessed the distribution of their penetration into the tissue domain. As expected, the tracer concentration (Fig. 3e-h, Fig. S4b) was lower in the regions directly above the bars of the mesh support. Fluid flow had a minimal impact on tracer distribution (Fig. 3f) and mean concentration (2.5% increase; Fig. 3g), even at high tissue permeability (1×10^−10^ m^2^). Uniformity increased at greater z planes further from the mesh and over time due to diffusion, evident by the 3-fold decrease in standard deviation (Fig. 3e). To test the predictions of distribution experimentally, we loaded a device with a slice of agarose embedded with biotinylated beads and injected a fluorescently-labeled soluble ligand (NeutrAvidin Rhodamine Red-X, or NRho) into the recirculating media.^37^ As predicted by the COMSOL model, the ligand reached the majority of the sample, with more protein delivered through the open regions of the mesh (Fig. 3g). In the future, area of the crossbars can be minimized or rearranged as needed, or customized membrane inserts may be used instead.

### Lymph node tissue slices remain viable for 24 hr culture under recirculating flow on-chip

Next, we tested for any impact of recirculating flow or co-culture on viability of tissues on the 3D printed chip, focusing on ex vivo lymph node slices because of the intended application to vaccination and because leukocytes are highly sensitive to their environment. Indeed, resin 3D-printed materials fabricated from commercially available resins are highly cytotoxic to primary murine splenocytes.^51,64^ However, we recently showed that coating the materials with a layer of parylene C, a method commonly used for medical implants and hardy cells in 3D-printed chips,^84–87^ was sufficient to protect splenocytes for at least 24 hr.^54^ Here, we confirmed that parylene C coating similarly restored viability of primary LN slices in static culture (Fig. 4a,b). Furthermore, LN slice viability was not significantly impacted by fluid flow on the device (Fig. 4a,c). Due to the heterogeneity of cell number across LN slices, there is a large variation in MTS signal in live samples as the assay is sensitive to number of cells present. We note that although not statistically significant, the high pump speed (75 µm/s) trended lower in viability than the other conditions, indicating possible damage that should be explored in the future. At the low speed (30 µm/s), viability was similar whether one or two slices were cultured in the device, and was not different between upstream and downstream culture wells (Fig. 4d). These results confirmed that tissue slices could be cultured in either well and in tandem while maintaining viability.

**Figure 4.**
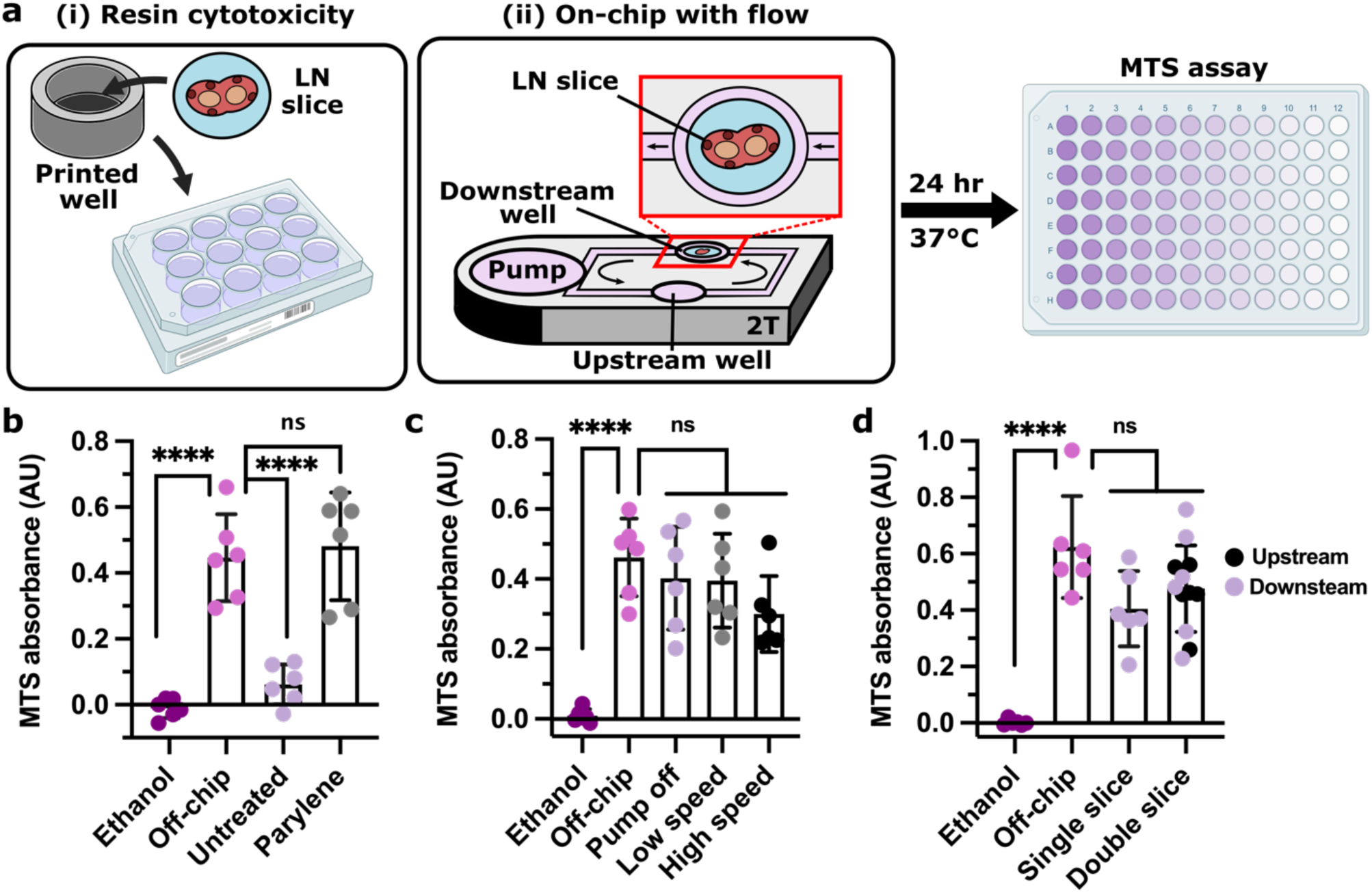
Lymph node slices were viable for 24 hr culture on-chip. (a) Experimental setup for 24 hr LN slice viability for (i) resin cytotoxicity and (ii) on-chip culture under fluid flow (well plate images created using BioRender.com). (b) MTS assay absorbance of LN slices cultured in untreated and parylene-coated 3D-printed wells (Clear resin) for 24 hrs, without fluid flow, compared to live (off-chip) and killed (ethanol) slices cultured off-chip. (c) MTS assay absorbance of LN slices cultured for 24 hrs on the parylene-coated 2T device (Clear resin) with the pump off, at low speed (1.2 V, 35 µm/s), or at high speed (1.7 V, 75 µm/s) compared to live (off-chip) and killed (ethanol) slices cultured off-chip. (d) MTS assay absorbance of LN slices cultured for 24 hrs on the parylene-coated 2T device (Clear resin) with one or two slices cultured per device at a low pump speed (1.2 V, 30 µm/s) compared to live (off-chip) and killed (ethanol) slices cultured off-chip. One-way ANOVA with Tukey post-hoc tests (n = 6). **** indicates p < 0.0001, ns indicates p > 0.1. Bars represent mean and standard deviation. Each dot represents one LN slice. All results pooled from two independent experiments.

### Two-compartment chip mimics antigen drainage to and processing in lymph node slices

Having established that the chip and pump provided recirculating fluid flow and molecular transport and was compatible with lymph node culture, we sought to use this multi-compartment system to model entry of molecular cues into the LN. *In vivo*, LNs are continually exposed to molecules arriving from upstream tissues, both directly through the draining lymphatics and indirectly via the bloodstream. We hypothesized that the multi-compartment chip could approximate the relative dilution that occurs after intravenous (i.v.) injection versus subcutaneous (s.c.) or intramuscular (i.m.) injection.^88^ To test this hypothesis, we loaded a biotinylated model tissue into one of the tissue wells and injected NRho into an upstream compartment that was connected either through the pump well (well first, approximating i.v. injection) or directly through a microchannel (channel first, approximating s.c. or i.m. injection) (Fig. 5a). For the first hour, the channel first condition had a nearly 10-fold greater rate of capture compared to the well first condition (Fig. 5b), resulting in a level of signal that it took the well-first condition 24 hr to attain. Thus, the device successfully replicated this aspect of the different drainage routes found *in vivo*. Because there was no protein clearance in this system, rates were similar after the first hour, and we anticipate that eventually, both conditions would reach equilibrium with comparable protein capture, unlike *in vivo* where molecules are gradually cleared from the body.

**Figure 5.**
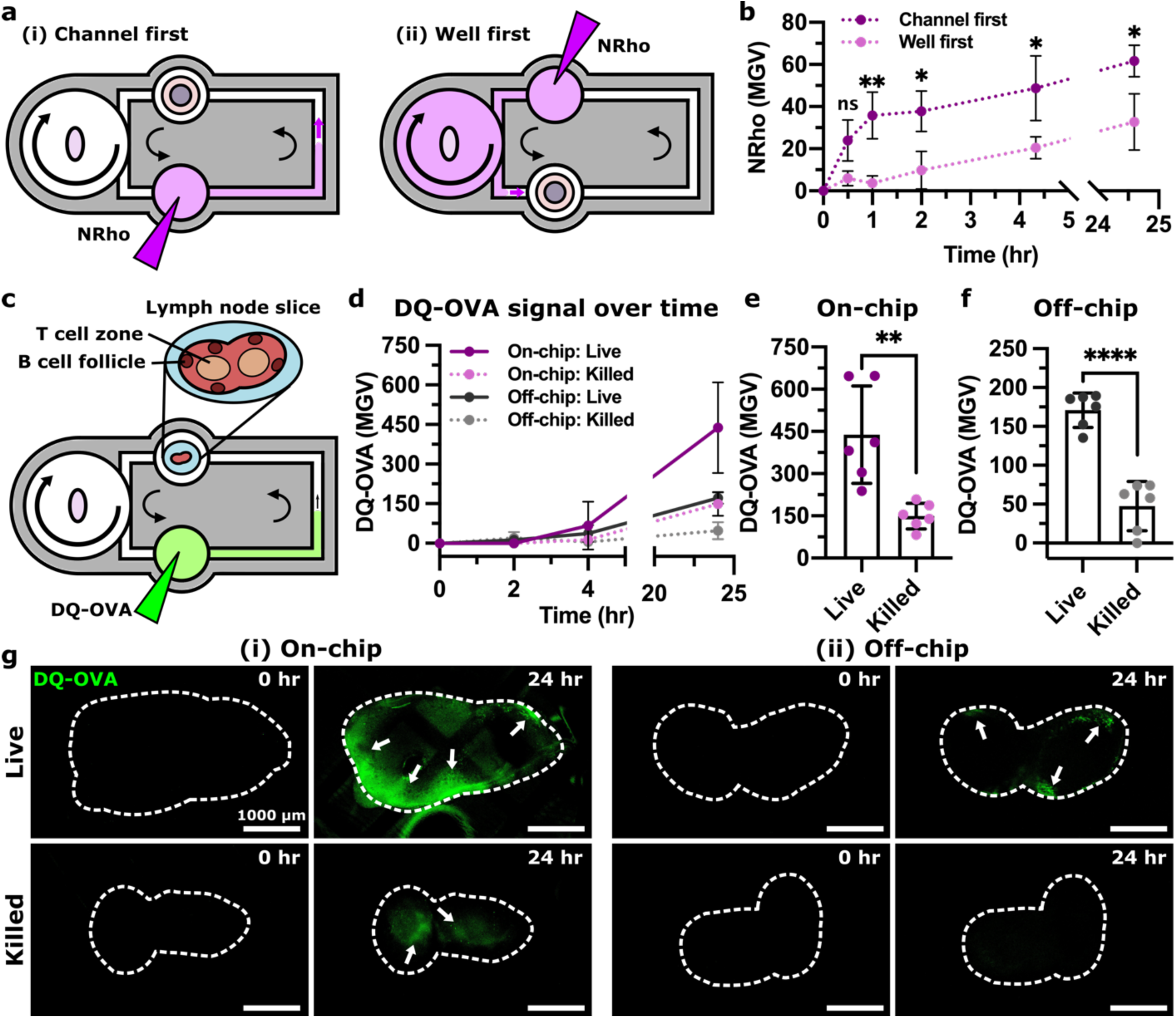
Modeling antigen drainage and uptake using two-compartment chip. (a) NRho (*purple*) was inserted in a filled 2T device (uncoated ITX-PEGDA resin) in the (i) channel first orientation and the (ii) well first orientation. (b) Quantification of NRho MGV in channel first and well first conditions over time (n = 3). One-way ANOVA with Tukey post-hoc tests. ns indicates p > 0.3, * indicates p < 0.03, and ** indicates p < 0.007. Dots and error bars represent mean and standard deviation, respectively. (c) A lymph node slice was inserted in a 2T device (parylene-coated Clear resin, 1.2 V, 30 µm/s) with a DQ-OVA injection (*green*) in the upstream culture well. (d) Mean gray value (MGV) of DQ-OVA in LN slices over time, showing processing of protein antigen. Dots and error bars represent mean and standard deviation; some error bars too small to see. (e,f) MGV of DQ-OVA in live or ethanol-treated LN slices cultured on-chip or off-chip at 24 hr. Unpaired t test (n = 6). **** indicates p < 0.0001, ** indicates p < 0.003. Each dot represents one LN slice. (g) Representative images of DQ-OVA signal (green) in live and killed slices cultured (i) on-chip and (ii) off-chip at 0 hr and 24 hr. Slices outlined with dashed white line from brightfield images (not shown). Arrows indicate regions that appear to have processed DQ-OVA. All results pooled from two independent experiments.

Next, we tested the ability of the multi-compartment system to model antigen drainage and processing in *ex vivo* LN slices. We selected DQ-ovalbumin (DQ-OVA) as a model antigen, as it becomes fluorescent only when proteolytically cleaved. DQ-OVA was injected into the upstream culture well to mimic s.c. injection (Fig. 5c) or added to the media in a well plate (off-chip) for comparison. As expected, the DQ-OVA signal increased at a greater rate in live LN slices than in ethanol-treated killed controls (Fig. 5d-g), both on-chip and off-chip, confirming that the live slices remained metabolically active and able to process antigen on-chip. A slow appearance of signal in ethanol-treated slices may have been due to residual protease activity (Fig. 5d);^89,90^ we observed similar results in formalin fixed tissues previously.^43^ We note that the DQ-OVA signal magnitude and distribution was variable between different lymph node slices, due to the heterogenous cell distributions found in the tissue. Processed antigen was brighter in slices on-chip than off-chip, consistent with better delivery by fluid flow than in static culture. After 24 hrs, the live slices cultured on-chip showed the mesh support pattern in certain regions (Fig. 5gi), similar to prior tests in mock tissue (Fig. 3g). Nevertheless, processed antigen appeared in similar regions of the lymph node as observed previously, near the outer sinus regions.^43,91^ Thus, the multi-compartment chip successfully modeled lymphatic drainage, phagocytosis, and processing of whole protein antigens in a lymph node.

### Acute immune response to on-chip vaccination was comparable to *in vivo* vaccination

Within minutes to hours of a vaccine injection, vaccine components drain from the site of injection to local LNs, where an immune response begins to develop.^28^ We sought to model these events by vaccinating a mock injection site upstream of a murine LN slice on the multi-compartment chip (Fig. 6a-i). As a model vaccine, we chose rhodamine-labeled ovalbumin (Rho-OVA) and R848 (TLR7/TLR8 agonist) as the antigen and adjuvant, respectively. This mixture, or a phosphate-buffered saline (PBS) control, was loaded into a block of soft hydrogel as the mock injection site. The calculated drainage time from the upstream culture well to the downstream culture well was approximately 23 min. To assess the accuracy of the multi-compartment chip and impeller pump platform, we benchmarked the on-chip response to vaccination against the response to s.c. vaccination *in vivo* in mice (Fig. 6a-ii). Furthermore, we compared the response on-chip to off-chip in static well plates to assess the utility of the chip versus conventional culture (Fig. 6a-iii).

**Figure 6.**
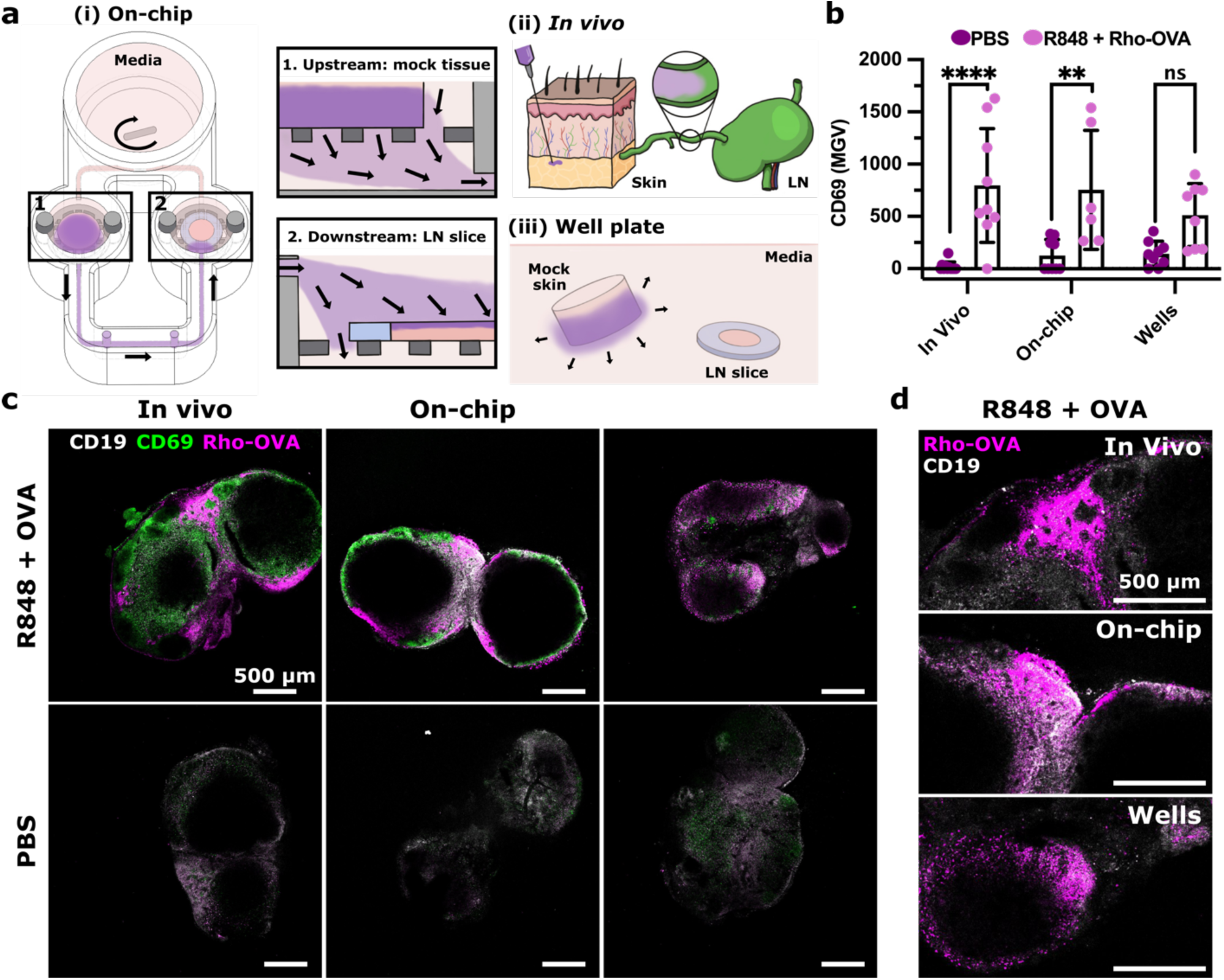
Similar activation marker signal and antigen distribution upon vaccination in *in vivo* and on-chip conditions. (a) Three vaccination conditions were compared: (i) on-chip in a 2T device (parylene-coated Clear resin), (ii) s.c. injection *in vivo*, and (iii) static well plate, where the vaccine is shown in purple. (b) Quantification of the MGV of CD69 in the CD19+ region for all conditions. Results in (e) were pooled from three independent experiments. Two-way ANOVA; **** indicates p < 0.0001, ** indicates p < 0.003, and ns indicates p > 0.07. Each dot represents a single LN slice. Bars represent standard deviation. (c) Representative images of LN slices with R848 + Rho-OVA or PBS from *in vivo* culture, on-chip culture, and well plate culture. B cells (CD19) are shown in grey; CD69 is shown in green; and Rho-OVA is shown in magenta. (d) Representative images of Rho-OVA (magenta) distribution across conditions. B cells (CD19) are shown in grey.

First, we assessed the early markers of activation after vaccination by confocal microscopy (Fig. S6), imaging three markers: CD69 for lymphocyte activation, and CD40 and CD86 for antigen-presenting cell (APC) activation. Because CD69 expression increases in LN slices after overnight culture,^92^ we restricted the *ex vivo* cultures to 6 hrs; *in vivo* vaccination was analyzed at a standard time point (24 hrs). Both on-chip and *in vivo* vaccinations both showed strong induction of CD69, particularly in the CD19+ region, whereas little response was observed in PBS controls (Fig. 6b,c). In contrast, the slices cultured off-chip yielded a smaller and not statistically significant increase in CD69 signal; however, it is not possible with the current data to conclude whether or not static conditions may have led to slower or lesser CD69 protein expression compared to conditions under flow. CD40 and CD86 both had high enough basal expression in PBS controls that vaccination had no impact; this result was consistent between *in vivo*, on-chip, and well plate conditions (Fig. S7). Despite the difference in time point, antigen distribution also was similar across all three platforms, with Rho-OVA appearing primarily in the sinus region (Fig. 6d).

Next, we compared transcriptomic changes in the LN by bulk RNA sequencing. For simplicity, we maintained the same 6 hr time point for all vaccination conditions (Fig. 7a). Principal component analysis (PCA) separated the vaccinated (R848 + Rho-OVA) and unvaccinated (PBS) conditions along PC2, although as a smaller effect than the separation between *in vivo* lymph nodes and *ex vivo* lymph node slices on-chip or in wells along PC1 (Fig. 7b). In future experiments, it would be preferable to slice the lymph nodes from the *in vivo* condition as well, as slicing likely impacted both gene expression and RNA yield. Nevertheless, heatmap analysis showed similar changes in gene expression after vaccination on-chip as *in vivo* or in well plates (Fig. 7c). Consistent with the earlier immunofluorescence staining (Fig. 6b), cd69 expression increased upon vaccination across all three conditions (Fig. 7di). Interestingly, cd40 expression also increased in all conditions (Fig. 7dii) and cd86 increased *in vivo* (Fig. S8), which were not detectable by immunofluorescence (Fig. S7b,c). As expected during an antiviral immune response,^93,94^ expression of genes such as ifng, tnf, and cxcl9 increased (Fig. 7diii-v) and decreased for il9r, a receptor for a cytokine commonly associated with an anti-inflammatory immune response (Fig. 7dvi).^95^ These changes were similar across all three conditions.

**Figure 7.**
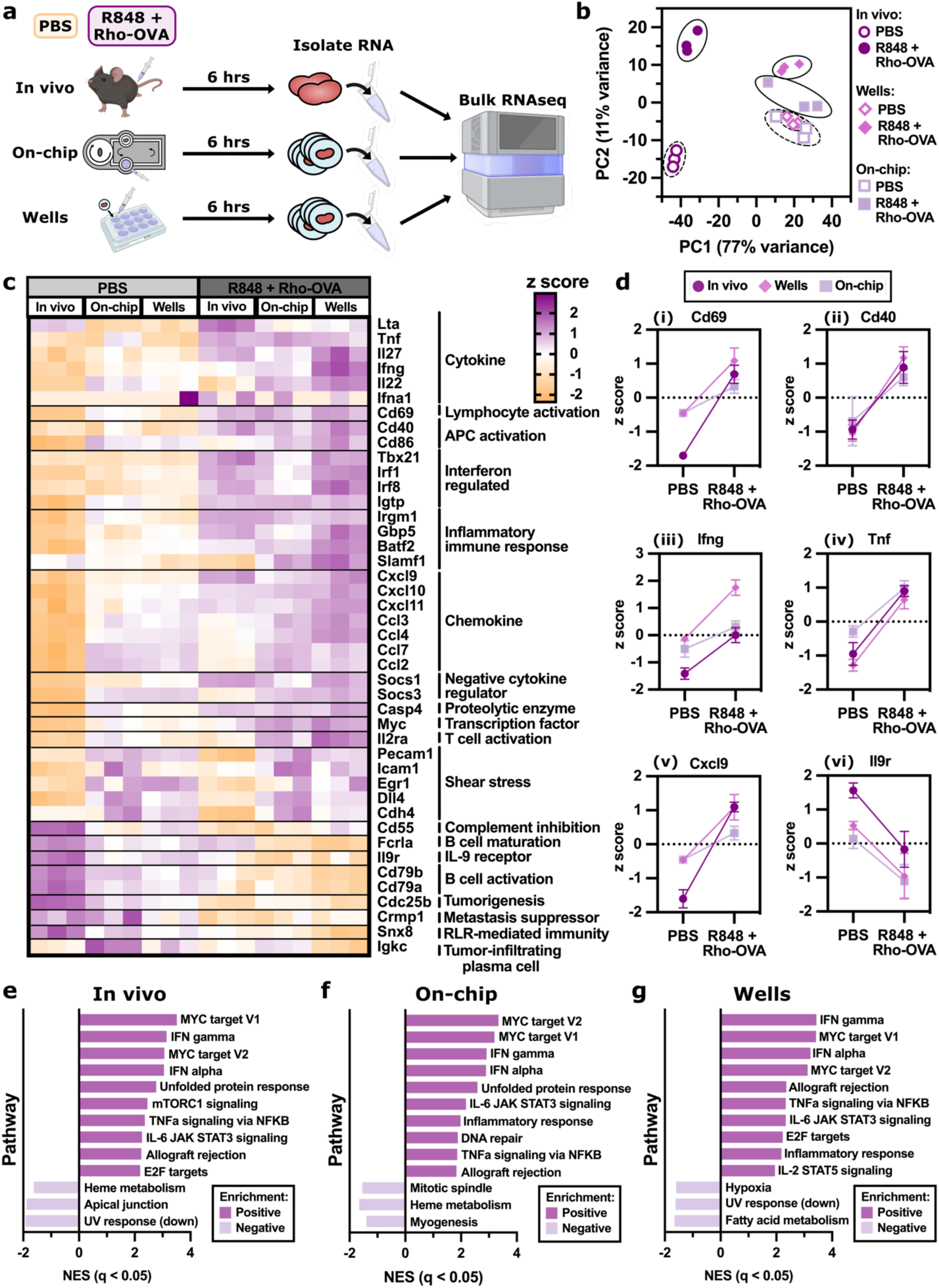
On-chip vaccination induced similar differentially expressed genes compared to *in vivo* vaccination. (a) Schematic showing RNA sequencing experimental workflow. For the *in vivo* condition, naive mice were injected s.c. with either PBS or R848 + Rho-OVA. After 6 hrs, the skin-draining lymph nodes were collected and the RNA was isolated from intact nodes before sequencing. For well and on-chip conditions, skin-draining lymph nodes from naive mice were first sliced then cultured *ex vivo* in a well plate or on the 2T device with either PBS or R848 + Rho-OVA. After 6 hrs, RNA was isolated from the tissue slices before sequencing. (b) Principal component analysis (PCA) of the normalized gene expression data, where the percent variance explained by each PC is listed in the axis labels. Colors and symbols represent different culture conditions (*in vivo*, on-chip, and wells) with either PBS and R848 + Rho-OVA. (c) Heatmap showing the expression of 36 selected genes of relevance to an immune response, where the cell value is the normalized z score. (d) The change in z score with the addition of R848 + Rho-OVA across *in vivo,* on-chip, and wells for (i) cd69, (ii) cd40, (iii) ifng, (iv) tnf, (v) cxcl9, and (vi) il9r. (e-g) Enriched pathways from the Hallmark gene set for (e) *in vivo*, (f) on-chip, and (g) wells comparing R848 + Rho-OVA to PBS (Table S3-5). The top 10 positively enriched pathways and the top 3 negatively enriched pathways are shown for each condition with q < 0.05.

Gene set enrichment analysis (GSEA) showed that both on-chip and well plate conditions shared eight of the top ten positively enriched pathways with the *in vivo* condition (Fig. 7e-g, Table S3-5). In fact, all conditions shared the same top four positively enriched pathways consistent with an early vaccine response: IFNγ response, IFNα response, MYC target V1, and MYC target V2. Of the top three negatively enriched pathways, on-chip and well plate conditions each shared one pathway with the *in vivo* condition.

Overall, the multi-compartment chip provided vaccine drainage and initial immune responses in the LN that were comparable to *in vivo* vaccination in terms of spatial distribution of antigen and activation markers as well as several of the most enriched gene expression pathways. Off-chip treatment of LN slices with this particular vaccine also produced a similar response, showing that in some cases, LN slices may be sufficient without the multi-organ culture platform. However, for vaccination conditions that require events to take place at a physically removed injection site, we expect that the multi-organ system will prove essential.

## CONCLUSIONS

Here, we have reported a user-friendly, 3D-printed multi-compartment chip and tubing-free impeller pump for the culture of one or more tissue models under biomimetic recirculating fluid flow. In this system, users may simply pipet media into the chip, load their tissues, and plug in a small, inexpensive control box, bypassing the need for peristaltic or syringe pumps. To replicate activity in spatially organized tissues such as the LN, we designed the device to support live *ex vivo* tissue slices, although the device is also compatible in principle with 3D cultures. By connecting murine or mock tissue slices using recirculating fluid flow, the multi-compartment chip modeled drainage of a vaccine from an upstream injection site to a downstream LN slice. On-chip vaccination yielded similar distribution of antigen, location and intensity of activation markers by immunofluorescence, and gene-specific expression patterns compared to *in vivo* vaccination.

Unlike static or mixed culture models, a strength of the 3D-printed chip and pump platform is it spatially compartmentalizes the events of an upstream organ compartment from those in a downstream compartment, with communication only via the media. As with other compartmentalized multi-organ chip systems, fluidic coupling of physically separated cultures will be important when modeling systemic effects of local events.^7,11,13,17^ In the context of lymph node communication, applications may include models of depot vaccination, tissue-specific infection, and local inflammation.

This multi-compartment chip and impeller pump are not without limitations. The current removable mesh geometry yielded little fluid flow through the tissue itself. For some applications, it will be useful to alter the geometry to reduce the gaps around the slice and increase the channel speed to incorporate the impacts of interstitial fluid flow.^15,80,96^ Furthermore, here we focused primarily on using flow for simple mass transfer between compartments; future work will be needed to address organ-to-organ scaling if required for particular applications.^97–101^ The volumes and channel lengths could be adjusted in the future to enable proper organ-to-organ scaling depending on the requirements of the modeled system.^7,97–101^ A benefit of the 3D printed fabrication is that the system is adaptable, so that the user can adjust geometries and features as needed, within the resolution of their printer. Currently, due to resolution limitations, the channel size for the multi-compartment chip was 500 µm, larger than devices fabricated using other methods such as soft lithography. As the resin 3D printing field continues to progress, higher resolution printers and smaller internal channel sizes may become routinely available.^53,67,68^ Alternatively, fabrication of the chip by injection molding or CNC machining would improve scalability and resolution. Finally, further work is needed to ensure that the rotation of the stir bar does not negatively impact recirculating lymphocytes, which was not required for the model developed here. Alternative impeller designs such as those used in cardiac impeller pumps may prove useful here.^102,103^ Furthermore, although we suspect that the parylene coating will retain biocompatibility for 7+ days based on prior reports,^85–87^ culture times longer than 24 hr will need to be confirmed with lymphocytes or LN slices.

While the current platform generated a similar short-term response to vaccine compared to *in vivo*, there are still many features of vaccination to expand on in future work. Currently, LN slices are best used for less than 24-48 hr, after which lymphocytes begin to egress, at least in static cultures.^43^ In the future, adjustments to culture conditions to enable longer culture times for LN slices, or incorporation of 3D cultured models instead of tissue slices, may enable models of longer-term responses to vaccination. Excitingly, the two-compartment system may enable incorporation of depot adjuvants and tissue-resident APCs into the upstream injection site, thus generating microphysiological models of clinically relevant vaccines such as those adjuvanted with aluminum salts. In the future, this platform can be used to model multi-tissue immunity by co-culturing a LN slice with additional upstream tissues. Beyond vaccination, we envision that this user-friendly, multi-compartment system will be compatible with additional tissue models to predict the progression of other complex phenomena including neurodegeneration, autoimmunity, and tumor immunity.

## AUTHOR CONTRIBUTIONS

SRC and RRP conceptualized the technology, planned experiments, interpreted data, and wrote the manuscript. SRC designed and fabricated the technology, performed the experiments, and analyzed the data. AGB contributed to the design and performance of vaccination experiments, including *in vivo* vaccination and related image analysis. AM led the analysis of RNAseq data through the Bioinformatics Core. All authors edited and approved the final manuscript.

## SUPPORTING INFORMATION

The Supplementary data file includes all supplemental figures, supplemental tables, and the caption for Movie S1.

## DATA AVAILABILITY

The data generated in this study, including source data for the figures and all device and pump design files are posted under Cook *et al.* “Replication Data for: A 3D-printed multi-compartment organ-on-chip platform with a tubing-free pump models communication with the lymph node,” at https://dataverse.lib.virginia.edu/dataverse/PompanoLab, available upon publication. The RNAseq data generated in this study is available through NCBI GEO using the accession GSE268137.

## ACKNOWLEDGEMENTS

The authors thank Kirk Bryson (Shared Materials Instrumentation Facility, Duke University) for his assistance and helpful correspondence. The authors thank Pankaj Kumar (Bioinformatics Core Facility, University of Virginia, RRID: SCR_012718) for his assistance with RNAseq data analysis and helpful correspondence. The authors thank Hannah Musgrove for technical assistance with cell culture preparation, printing devices, and running initial viability experiments. The authors thank Geane Miranda for technical assistance with printing devices and pump assembly, Alyssa Montalbine for technical assistance with printing devices, Erin Lawrence for technical assistance with LN slicing and editing support, and Meredith Harris for editing support.

Research reported in this publication was supported by the National Institute of Allergy and Infectious Diseases of the National Institute of Health under Award Number R01AI174207 and R01AI131723. The content is solely the responsibility of the authors and does not necessarily represent the official views of the National Institutes of Health. SRC was supported in part by the 2021 Presidential Fellowship for Collaborative Neuroscience from the UVA Brain Institute, the 2022 Sidney M. Hecht Graduate Fellowship from UVA Chemistry, and a 2021 Research Grant from UVA Graduate School of Arts and Sciences Committee (GSASC). Parylene coating was performed at the Duke University Shared Materials Instrumentation Facility, a member of the North Carolina Research Triangle Nanotechnology Network, which is supported by the National Science Foundation (Award ECCS-2025064) as part of the National Nanotechnology Coordinated Infrastructure.

