## Supplemental information for "A 3D-printed multi-compartment organ-on-chip platform with a tubing-free pump models communication with the lymph node"

Electronic Supplementary Information for


**Contents:**


- Supplemental Methods
- Supplemental Movie Caption
- Supplemental Figures
- Supplemental Tables

### I. SUPPLEMENTAL METHODS

#### Characterization of fluid flow and shear stress on-chip using COMSOL Multiphysics

The fluid flow profile through the tissue culture well was modeled in three dimensions using the free and porous media flow module and transport of diluted species in porous media module of COMSOL Multiphysics (Version 6.1). The computational model matched the 3D geometry of the tissue culture well. The culture chamber was split into two cylinders, where the bottom chamber had a diameter of 6.2 mm with a height of 1.5 mm, and the top chamber had a diameter of 7.7 mm with a height of 2.5 mm. An inlet channel was connected to the top of the top culture well, and an outlet channel was connected to the base of the bottom culture well on the opposite side. Both channels had 0.5 x 0.5 mm cross-section with a length of 15 mm. The tissue slice was modeled as a cylinder with a diameter of 5 mm and a height of 0.3 mm, within which the inner 3 mm represented tissue and a surrounding 1 mm ring represented 6% agarose. Aqueous media was modeled as an incompressible fluid with a viscosity of 1.00 mPa s and a density of 1000 kg/m$^3$. The tissue was modeled as a porous matrix with a viscosity of 1.00 mPa s, a density of 1000 kg/m$^3$, a porosity of 0.2, and a permeability ranging from 1x10$^{-10}$ m$^2$ to 1x10$^{-12}$ m$^2$.[25,50] The 6% agarose was modeled as a porous matrix with a viscosity of 1.00 mPa s, a density of 1000 kg/m$^3$, porosity of 0.2, and a permeability of 4.26x10$^{-18}$ m$^2$.[51] The mesh support geometry was excluded from the physics to generate a wall around the geometry. A "normal" triangular mesh was used as generated by the software. The inlet velocity was set to a maximum velocity of 30 µm/s within the channel, unless stated otherwise. The outlet was set to atmospheric pressure. The simulation was solved in time-dependent mode, and the readouts were reported at 5 min after reaching steady state unless noted otherwise. The velocity and shear stress through the tissue were measured along cut lines through the center of the slice (z = 0.15), 10 µm from the top (z = 0.29), and 10 µm from the bottom (z = 0.01). Unless otherwise noted, the central cut line was used. When testing different inlet speeds, the inlet velocity was set to a maximum velocity of 10 μm/s, 20 μm/s, 30 μm/s, 50 μm/s, 75 μm/s, 100 μm/s, and 150 μm/s within the channels.

The computational model was used to predict the fluid shear stress (FSS) and shear rate at the channel wall and the central tissue cutline. The inlet velocity was set to the range of maximum velocities listed above. The FSS (dyn/cm$^2$) was approximated using Equation (1), where $\gamma$ is shear rate (1/s) and $\eta$ is viscosity (Pa s).

$$FSS = 0.1\gamma\eta \tag{1}$$

#### II. SUPPLEMENTAL MOVIE CAPTION

**Movie S1**. A time-lapse video of recirculating fluid flow on a two-tissue device (Clear resin) with an agarose slice in each well, where blue dye was inserted into the upstream culture well (Figure 2d). The time was displayed in hr, min, and sec on the timer. The pump well inner diameter was 15 mm for scale.

#### III. SUPPLEMENTAL FIGURES

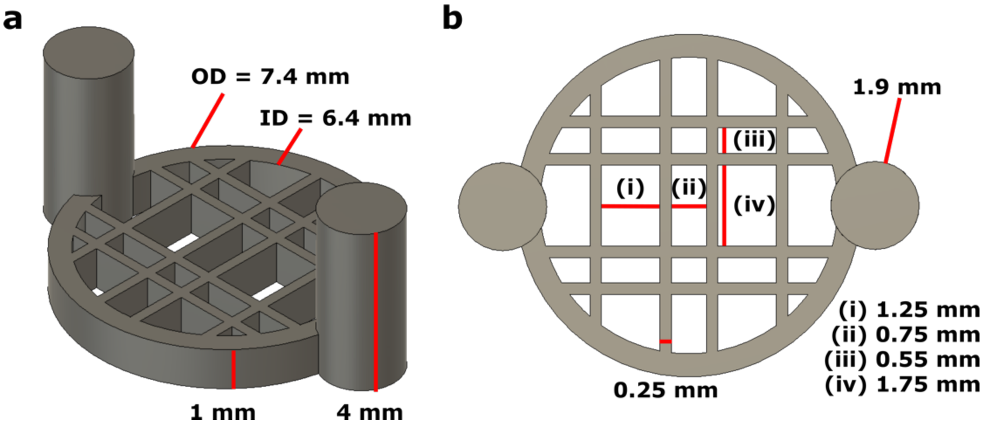


**Figure S1. Dimensions of removable mesh insert**. 3D renderings of the insert viewed (a) from the side and (b) from the top. An angled side view of the removable mesh insert.

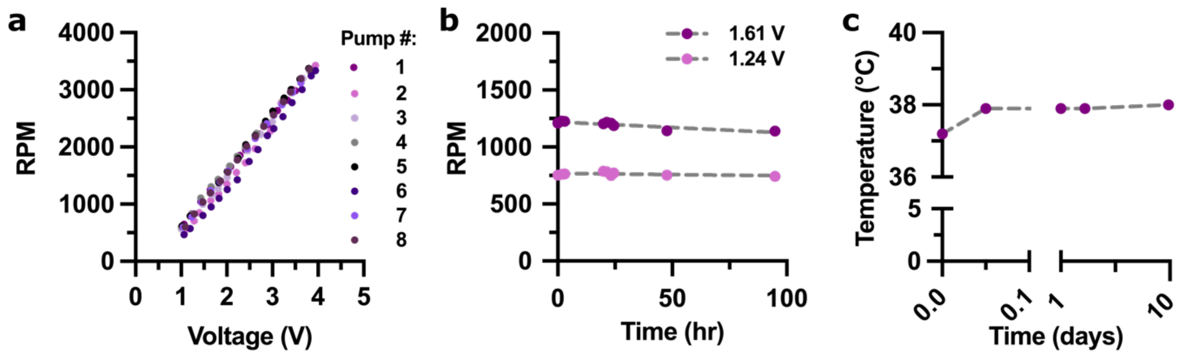


**Figure S2. Motor-based impeller pump characterization.** (a) As the voltage increased, the RPMs for each pump increased linearly, with little variation between pumps. (b) The RPM stability at two different pump voltages over a 90 hr time period. (c) The temperature within a cell culture incubator over 10 days with eight impeller pumps running at 4 V.

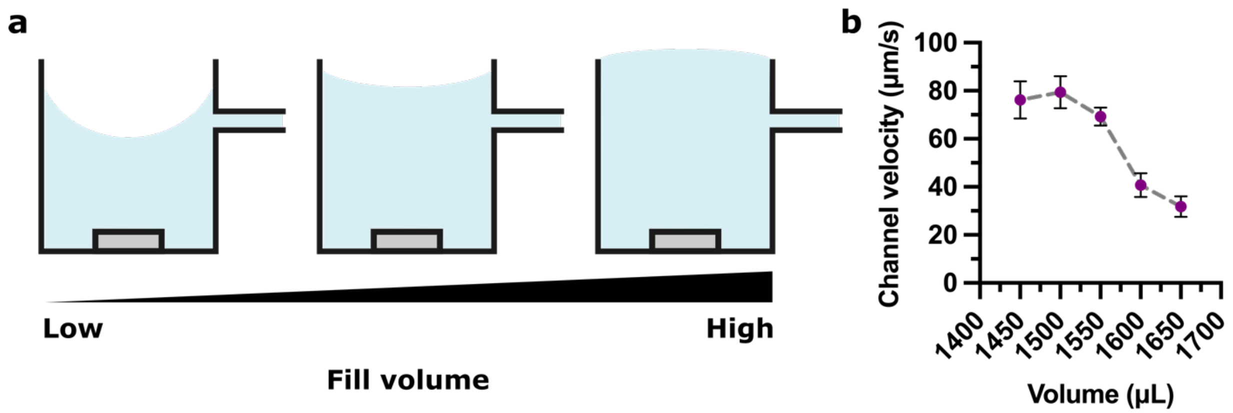


**Figure S3. Impact of fill volume on channel velocity.** (a) Schematic of the side view of the pump well and intersecting channel showing the impact of fill volume (*blue*) on the pump vortex. (b) Maximum velocity within the channel as a function of total fill volume, measured using a 5 mm stir bar at constant RPM (1.8 V, 1460 RPM) in a 0T device (no culture wells). Fill volumes ranging from low (1450 µL) to high (1650 µL) were tested.

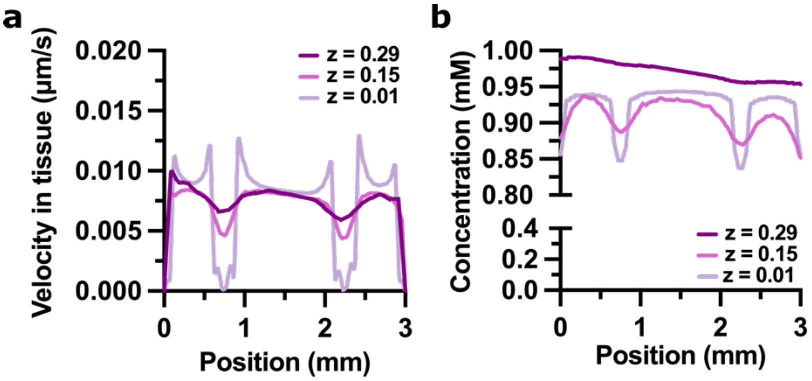


**Figure S4. Predicted velocity and tracer concentration in the tissue slice from COMSOL computational modeling.** Predicted (a) velocity and (b) protein concentration in tissue with a permeability of $1x10^{-10}$ $m^2$ at different z planes at t = 100 min with an inlet speed = 40 μm/s, where z = 0 mm is the bottom of the tissue slice and z = 0.3 mm is the top of the tissue.

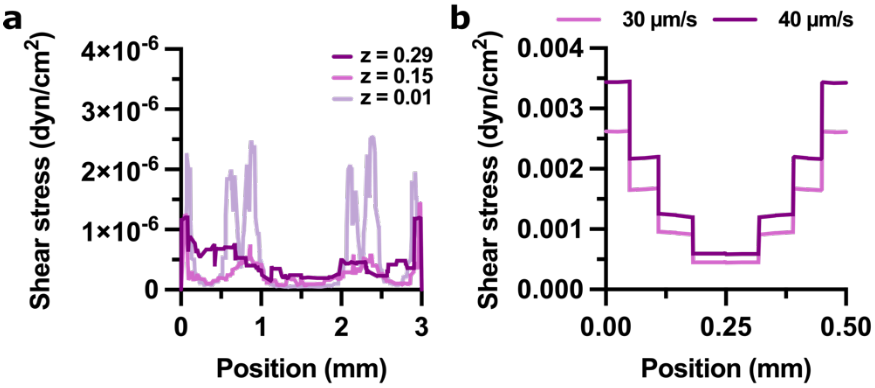


**Figure S5. Predicted shear stress using COMSOL computational modeling.** (a) Predicted shear stress through tissue at an inlet speed of 40 μm/s with a tissue permeability of $1x10^{10}$ $m^2$. (b) Predicted shear stress in the inlet channel at an inlet speed of 30 and 40 μm/s.

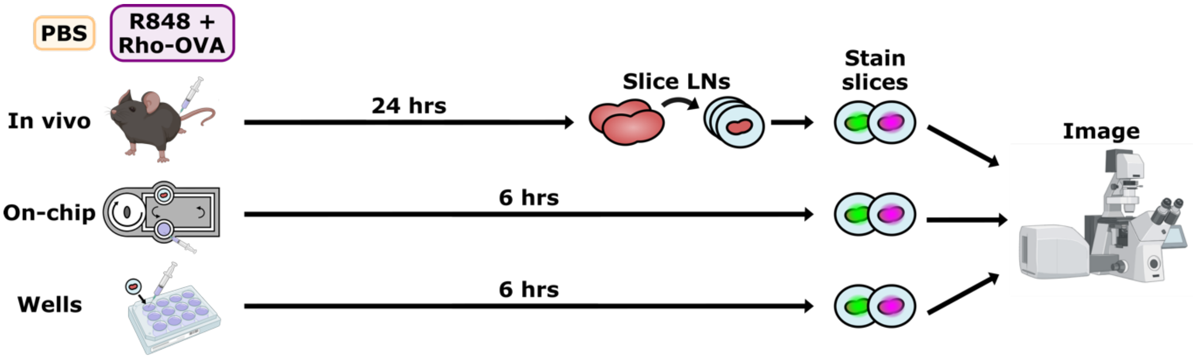


**Figure S6. Workflow for imaging portion of comparative vaccination experiment.** For the *in vivo* condition, naive mice were injected with either PBS or R848 + Rho-OVA. After 24 hrs, the skin-draining lymph nodes were collected, sliced, stained, and imaged. For well and on-chip conditions, skin-draining lymph nodes from naive mice were first sliced then placed in a well plate or on the device with either PBS or R848 + Rho-OVA. After 6 hrs, the slices were stained and imaged.

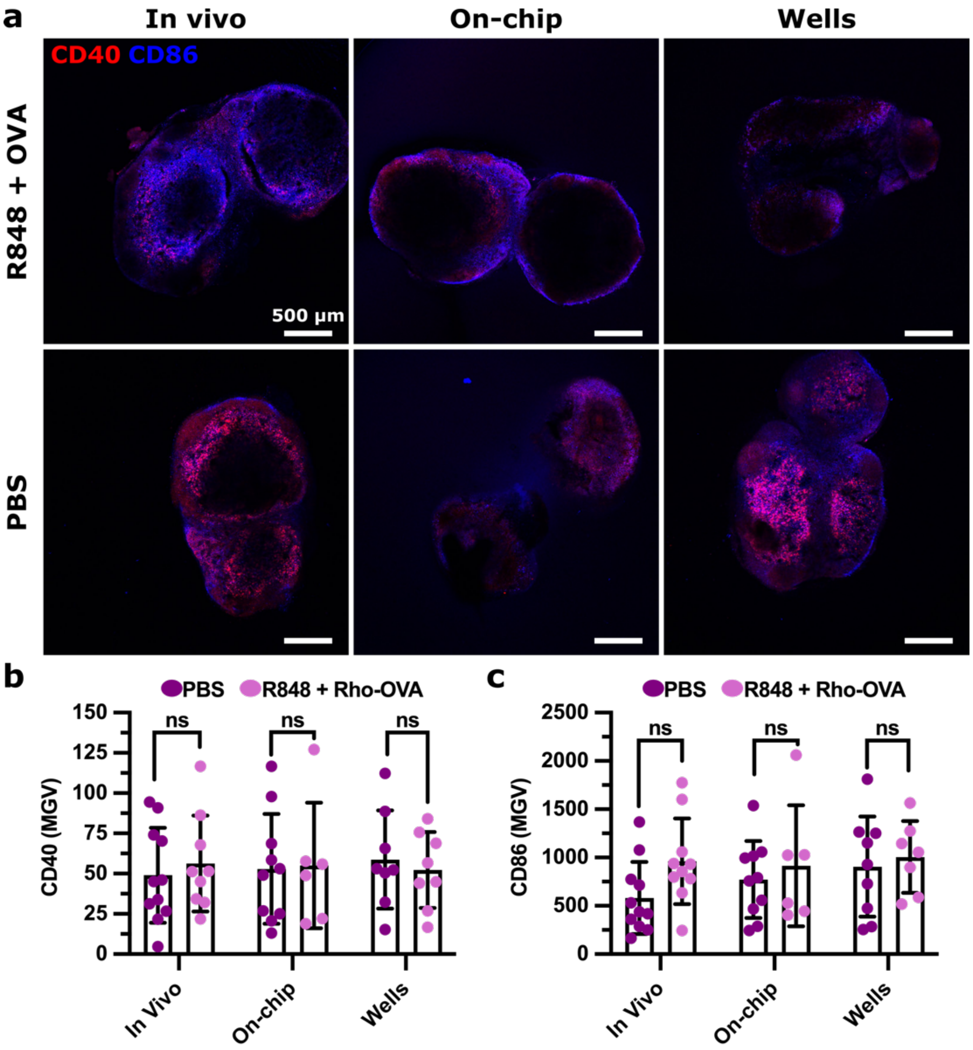


**Figure S7. No change in APC activation markers CD40 and CD86 signal upon vaccination.** (a) Representative images of lymph node slices with R848 + OVA and PBS only from *in vivo* culture (24 hr), on-chip culture (6 hr), and well plate culture (6 hr). CD40 is shown in red and CD86 is shown in blue. These images are from the same lymph node slices as shown in Figure 5, to enable comparison with the other markers shown in that figure. (a, b) Quantification of the MGV of (b) CD40 and (c) CD86 across the whole slice. Results were pooled from three independent experiments. Two-way ANOVA; ns indicates $p > 0.07$. Each dot represents a single LN slice. Bars represent standard deviation.

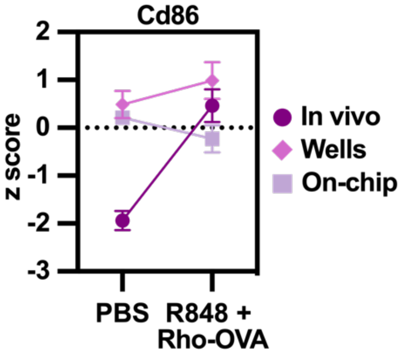


**Figure S8. Changes in gene expression for cd86.** The change in z score with the addition of R848 + Rho-OVA across *in vivo*, on-chip, and wells for cd86.

#### IV. SUPPLEMENTAL TABLES

**Table S1. Antibody cocktail used for comparative vaccination experiment.**

| *Target* | *Clone* | *Fluorophore* | *Product Number* | *Lot Number* | *Vendor* |
|---|---|---|---|---|---|
| CD16/32 | 93 | N/A | 101302 | B366439 | Biolegend |
| CD86 | GL-1 | Brilliant Violet 421 | 105032 | B364400 | Biolegend |
| CD69 | H1.2F3 | Alexa Fluor 488 | 104516 | B371402 | Biolegend |
| CD40 | 3/23 | Alexa Fluor 647 | 124614 | B380619 | Biolegend |
| CD19 | 6D5 | Starbright Violet 670 | MCA1439SBV670 | 100004959 | Bio-Rad |

**Table S2. Sample key for RNAseq experiment**

| *Sample number* | *Condition* |
|---|---|
| 5 | In vivo PBS |
| 11 | |
| 17 | |
| 1 | On-chip PBS |
| 13 | |
| 19 | |
| 3 | Wells PBS |
| 15 | |
| 21 | |
| 6 | In vivo R848 + Rho-OVA |
| 12 | |
| 18 | |
| 2 | On-chip R848 + Rho-OVA |
| 14 | |
| 20 | |
| 4 | Wells R848 + Rho-OVA |
| 16 | |
| 22 | |

**Table S1. Hallmark pathway norm. enrichment score (NES) for *in vivo* condition (q < 0.05)**

| *Rank* | *Pathway* | *Adjusted p value (q)* | *NES* |
|---|---|---|---|
| 1 | MYC targets V1 | 0.00549 | 3.50883 |
| 2 | Interferon gamma response | 0.00549 | 3.14648 |
| 3 | MYC targets V2 | 0.00549 | 3.04768 |
| 4 | Interferon alpha response | 0.00549 | 3.04783 |
| 5 | Unfolded protein response | 0.00549 | 2.76748 |
| 6 | mTORC1 signaling | 0.00549 | 2.45571 |
| 7 | TNFa signaling via NFKB | 0.00549 | 2.36461 |
| 8 | IL-6 JAK STAT3 signaling | 0.00549 | 2.26195 |
| 9 | Allograft rejection | 0.00549 | 2.23425 |
| 10 | E2F targets | 0.00549 | 2.19491 |
| 11 | Inflammatory response | 0.00549 | 2.19395 |
| 12 | Oxidative phosphorylation | 0.00549 | 2.06327 |
| 13 | G2M checkpoint | 0.00549 | 2.04077 |
| 14 | DNA repair | 0.00549 | 2.03129 |
| 15 | IL2 STAT5 signaling | 0.00549 | 1.84604 |
| 16 | UV response UP | 0.00549 | 1.81175 |
| 17 | Heme metabolism | 0.00664 | -1.62310 |
| 18 | Epithelial mesenchymal transition | 0.00664 | -1.81818 |
| 19 | Apical junction | 0.00664 | -1.88665 |
| 20 | UV response DN | 0.00664 | -1.91892 |
| 21 | Myogenesis | 0.00664 | -2.07587 |
| 22 | Bile acid metabolism | 0.00664 | -2.19790 |
| 23 | Mitotic spindle | 0.01224 | -1.40881 |
| 24 | Peroxisome | 0.01547 | -1.56988 |
| 25 | Apoptosis | 0.01579 | 1.46723 |
| 26 | Complement | 0.02039 | 1.41896 |
| 27 | Xenobiotic metabolism | 0.02098 | -1.39161 |
| 28 | P53 signaling DN | 0.02730 | 1.38355 |
| 29 | KRAS signaling DN | 0.02754 | -1.39789 |
| 30 | Fatty acid metabolism | 0.02754 | -1.43084 |
| 31 | P13K AKT mTOR signaling | 0.02834 | 1.44779 |

**Table S2. Hallmark pathway NES for on-chip condition (q < 0.05)**

| *Rank* | *Pathway* | *Adjusted p value (q)* | *NES* |
|---|---|---|---|
| 1 | MYC targets V2 | 0.00706 | 3.34360 |
| 2 | MYC targets V1 | 0.00706 | 3.19705 |
| 3 | Interferon gamma response | 0.00706 | 2.91494 |
| 4 | Interferon alpha response | 0.00706 | 2.89706 |
| 5 | Unfolded protein response | 0.00706 | 2.59232 |
| 6 | IL-6 JAK STAT3 signaling | 0.00706 | 2.174622 |
| 7 | Inflammatory response | 0.00706 | 1.98224 |
| 8 | DNA repair | 0.00706 | 1.88388 |
| 9 | TNFa signaling via NFKB | 0.00706 | 1.86379 |
| 10 | Allograft rejection | 0.00706 | 1.83635 |
| 11 | E2F targets | 0.00706 | 1.74667 |
| 12 | mTORC1 signaling | 0.00706 | 1.67963 |
| 13 | Oxidative phosphorylation | 0.00706 | 1.60907 |
| 14 | Mitotic spindle | 0.00706 | -1.54289 |
| 15 | Heme metabolism | 0.00706 | -1.65257 |
| 16 | Myogenesis | 0.01986 | -1.39735 |

**Table S3. Hallmark pathway NES for well plate condition (q < 0.05)**

| *Rank* | *Pathway* | *Adjusted p value (q)* | *NES* |
|---|---|---|---|
| 1 | Interferon gamma response | 0.00528 | 3.43666 |
| 2 | MYC targets V1 | 0.00528 | 3.42381 |
| 3 | Interferon alpha response | 0.00528 | 3.22391 |
| 4 | MYC targets V2 | 0.00528 | 3.12163 |
| 5 | Allograft rejection | 0.00528 | 2.35705 |
| 6 | TNFa signaling via NFKB | 0.00528 | 2.33754 |
| 7 | IL-6 JAK STAT3 signaling | 0.00528 | 2.33675 |
| 8 | E2F targets | 0.00528 | 2.22874 |
| 9 | Inflammatory response | 0.00528 | 2.18648 |
| 10 | IL-2 STAT5 signaling | 0.00528 | 1.95186 |
| 11 | G2M checkpoint | 0.00528 | 1.93762 |
| 12 | Unfolded protein response | 0.00528 | 1.75330 |
| 13 | DNA repair | 0.00528 | 1.71872 |
| 14 | Apoptosis | 0.00528 | 1.66961 |
| 15 | mTORC1 signaling | 0.00528 | 1.56138 |
| 16 | Oxidative phosphorylation | 0.00528 | 1.53443 |
| 17 | Hypoxia | 0.00714 | -1.59489 |
| 18 | UV response DN | 0.00714 | -1.60992 |
| 19 | Fatty acid metabolism | 0.00714 | -1.64992 |
| 20 | Bile acid metabolism | 0.00714 | -1.73781 |
| 21 | Epithelial mesenchymal transition | 0.00714 | -1.75378 |
| 22 | Myogenesis | 0.00714 | -2.28017 |
| 23 | Complement | 0.00963 | 1.50923 |
| 24 | Heme metabolism | 0.01286 | -1.53568 |
| 25 | Xenobiotic metabolism | 0.01834 | -1.47250 |
| 26 | Glycolysis | 0.02366 | -1.36421 |
| 27 | Peroxisome | 0.03429 | -1.47662 |
| 28 | UV response UP | 0.04530 | 1.38054 |
